# DNA nicks in both leading and lagging strand templates can trigger break-induced replication

**DOI:** 10.1101/2023.12.22.573028

**Authors:** Yuanlin Xu, Yassine Laksir, Carl A. Morrow, Kezia Taylor, Costas Tsiappourdhi, Patrick Collins, Su Jia, Christos Andreadis, Matthew C. Whitby

**Affiliations:** Department of Biochemistry, University of Oxford, South Parks Road, Oxford, OX1 3QU UK

## Abstract

Encounters between replication forks and unrepaired single-strand DNA breaks (SSBs) generate single-ended double-strand breaks (seDSBs) that can later become double-ended (deDSBs) through fork convergence. seDSBs can be repaired by break-induced replication (BIR), which is a highly mutagenic pathway that is thought to be repsonsible for many of the mutations and genome rearrangements that drive cancer development. However, the frequency of BIR’s deployment and its ability to be triggered by both leading and lagging template strand SSBs was unclear. Using site- and strand-specific SSBs generated by nicking enzymes, including CRISPR-Cas9n, we demonstrate that leading and lagging template strand SSBs in fission yeast are typically converted into deDSBs that are repaired primarily by error-free homologous recombination. However, both types of SSB can also trigger BIR, and the frequency of these events increases when the converging fork is delayed and the non-homologous end joining protein Ku70 is deleted.

## Introduction

DNA single-strand breaks (SSBs) are amongst the most pervasive DNA lesions found in cells, occurring at a frequency of 10^4^ - 10^5^ per cell per day^1^. These breaks can result from ionizing radiation and reactive oxygen species, and are formed as intermediates in several DNA metabolic processes, including base excision repair and DNA torsional stress relief by topoisomerase I (Top1)^1,2^. Because SSBs are so numerous, they can sometimes be encountered by DNA replication forks before they are repaired^1,2^. These encounters can cause genome instability and cytotoxicity, and also form the basis of cancer treatments that use Poly (ADP-ribose) polymerase (PARP) inhibitors and derivatives of the Top1 poison camptothecin (CPT) to increase SSB accumulation in cells^3–5^.

Insight into what happens when a replication fork encounters an SSB has in part come from studies of CPT-induced breaks. CPT acts by inhibiting the re-ligation step in Top1’s catalytic cycle, which results in the formation of a cleavage complex (Top1cc), consisting of a SSB with Top1 covalently attached at its 3’ end^3^. Initial studies showed that CPT could give rise to replication-associated DNA double-strand breaks (DSBs)^6–8^. It was later demonstrated that DSBs arose through a process of replication run-off where the replicative helicase CDC45-MCM2-7-GINS (CMG) translocates off the DNA at Top1ccs in the leading strand template^9^. Studies that employed DNA nicking enzymes to generate site-specific SSBs also showed that SSBs are converted into either single-ended (se) or double-ended (de) DSBs during DNA replication^10–14^. Most recently, biochemical experiments, using Xenopus egg extract, confirmed the generation of seDSBs by replication run-off at leading strand template SSBs (lead-SSBs)^15^. Surprisingly, seDSBs were also observed when the SSB was in the lagging template strand (lag-SSBs)^15^. However, unlike at lead- SSBs, where CMG runs off the DNA at the break, at lag-SSBs CMG was proposed to translocate along double-stranded DNA (dsDNA) beyond the break, and was then actively unloaded from the DNA by a ubiquitin-mediated process^15^.

Only a few studies have investigated the repair of DSBs that arise from replication run-off at site-specific SSBs and, most of those that have, analysed the repair of plasmid substrates where the opposing replication fork converges with the already broken fork to form a deDSB^13,16,17^. A key conclusion from these studies is that deDSBs that derive from replication run-off and fork convergence are predominantly repaired by homologous recombination (HR) through a process of equal sister chromatid exchange. Key steps in this process include: 1) nucleolytic resection of the break to generate 3’-ended single-strand DNA (ssDNA) tails; 2) nucleation of Rad51 onto the ssDNA mediated by Rad52 (in yeast) or BRCA2 (in humans); 3) spreading of Rad51 along the DNA to form a nucleoprotein filament; 4) a Rad51-mediated search for a homologous dsDNA; and 5) invasion of the nucleoprotein filament into the located homologous dsDNA to establish a displacement (D) loop at which new DNA synthesis is primed. Following these initial steps, repair can then proceed via synthesis-dependent strand annealing (SDSA) or through the formation and subsequent resolution/dissolution of a double Holliday junction if the DSB is double-ended^18–20^. However, if the break is single-ended, a different HR mechanism comes into play called break-induced replication (BIR).

BIR proceeds via continued elongation of the invading strand by a DNA polymerase coupled with migration of the D-loop^21,22^. This non-canonical form of DNA replication can copy an entire chromosome arm but is highly mutagenic and prone to template switching, generating various genome rearrangements that are thought to drive the development of genetic diseases including cancer^21–24^. Given its potential to disrupt genomes and cause disease, it remains uncertain whether broken replication forks are typically repaired through BIR. Indeed, BIR may be reserved for specific circumstances where alternative repair mechanisms and means for completing DNA replication are unavailable, such as when cells experience oncogene-induced replication stress or enter mitosis with unreplicated DNA^25–29^.

To date, only one study has directly investigated whether BIR is induced by replication run-off at SSBs, which was conducted in budding yeast and demonstrated that lead-SSBs can indeed cause BIR^14^. However, it remains unknown whether BIR is a common and conserved response to replication run-off, whether the nature of the SSB has an impact on pathway choice, and whether lag-SSBs can induce BIR. In this study, we investigate the ability of both lead- and lag-SSBs to induce BIR in the fission yeast *Schizosaccharomyces pombe*, which, in evolutionary terms, is about as distantly related to budding yeast as each yeast is to humans^30^. Using site- and strand-specific SSBs generated by CRISPR/Cas9 nickase (Cas9n), we demonstrate that most SSBs are converted into deDSBs that are repaired through Rad51-mediated sister chromatid recombination. However, we also find evidence that BIR can be induced from both lead- and lag-SSBs, and show that the propensity for BIR increases when fork convergence is delayed and the non-homologous end joining (NHEJ) protein Ku70 is deleted.

## Results

### Cas9n-induced SSBs are converted into deDSBs that strongly induce direct repeat recombination

To investigate the ability of lead- and lag-SSBs to generate DSBs and stimulate recombination, we designed a guide RNA (gRNA) to direct Cas9n to make a SSB in the DNA between two *ade6^-^* mutant heteroalleles on chromosome 3 of *S. pombe*. At this chromosomal site, DNA replication proceeds almost entirely in the telomere to centromere direction (Figure 1A). We employed two Cas9n variants, D10A and H840A, to introduce strand specific nicks. In our experimental system, the gRNA binds to the lagging strand template meaning that Cas9n^H840A^ generates a lead-SSB, whereas Cas9n^D10A^ generates a lag-SSB. To determine whether these SSBs give rise to DSBs, we examined genomic DNA extracted from asynchronously growing cell cultures using native gel electrophoresis. Southern blot analysis was then used to identify a ∼4.1 kb PstI-NheI restriction fragment that encompasses the gRNA binding site (Figure 1A and 1B). As anticipated, in cells expressing gRNA along with the fully active form of Cas9, we detected DNA bands of ∼2 kb in size using probes that hybridize to either side of the gRNA binding site (Figure 1B). The percentage of both DSB signals was similar, consistent with them corresponding to the two sides of a deDSB (Figure 1C). Reassuringly, no DSB signal was detected in genomic DNA from cells expressing the catalytically dead form of Cas9 (Cas9d) that contains both D10A and H840A mutations (Figure 1B and 1C). However, surprisingly, genomic DNA from cells expressing either Cas9n^H840A^ or Cas9n^D10A^, together with gRNA, displayed similar levels of deDSB signals as observed with Cas9 (Figure 1B and 1C).

**Figure 1.**
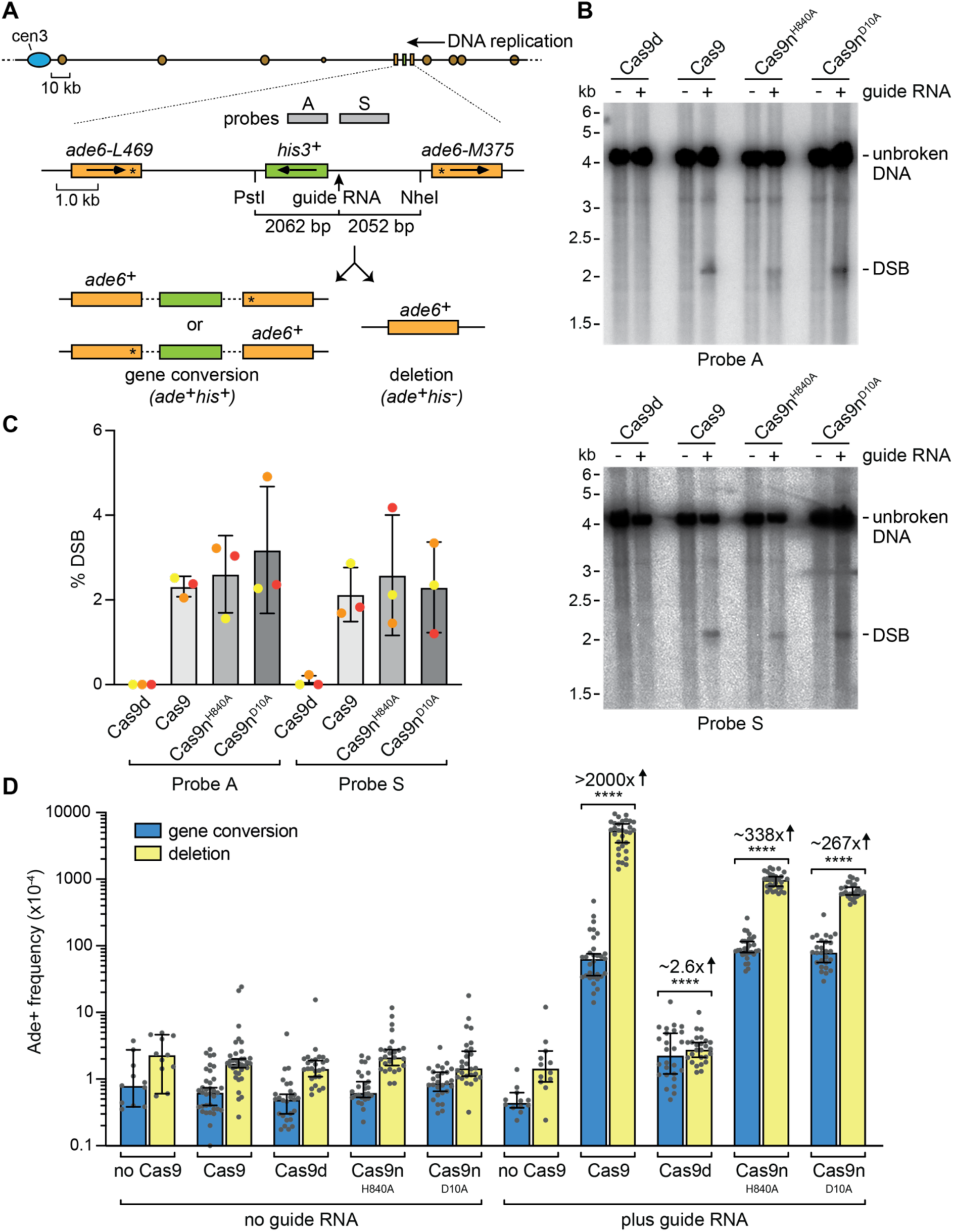
Cas9n-induced SSBs cause deDSBs that are highly recombinogenic. **(A)** Diagram showing the location of the gRNA binding site on chromosome 3. Replication origins (brown circles), probes (grey rectangles) and relevant genes (orange and green rectangles with arrows indicating the direction of transcription) are indicated. Asterisks indicate the position of loss of function mutations in the two *ade6* alleles. The two types of Ade+ recombinant are shown in the bottom panel. **(B)** Detection of DSBs in genomic DNA by Southern blot analysis. DNA was extracted from cells expressing the indicated version of Cas9 with and without gRNA. The DNA was cut with PstI and NheI, and the relevant restriction fragment was detected using probes A and S. **(C)** Quantification of the DSB signals detected by Southern blot analysis. Data are presented as mean values ± SD. Individual data points are shown as coloured dots, with the colour denoting which experiment the data are from. **(D)** Frequency of Ade+ recombinants in strains containing the *ade6^-^* direct repeat substrate shown in A. The presence of gRNA and version of Cas9 is indicated. Data are presented as median values ± 95% confidence interval with individual data points shown as grey dots. Fold changes are for total Ade+ frequencies and are relative to the equivalent no gRNA control strain. *P*-values are for the comparison to the equivalent no gRNA control strain and were calculated by the Mann-Whitney test (two-tailed). **** *p*-value <0.0001. The data are also reported in Supplementary Table 1, which includes the strain numbers, the number of colonies tested for each strain (*n*) and relevant *p*-values.

To measure the recombination induced by Cas9 and Cas9n, we monitored the frequency of Ade+ recombinants that arise from gene conversion and deletion recombination events between the two different copies of *ade6^-^*(Figure 1A and 1D). All three nuclease active forms of Cas9 induced high levels of direct repeat recombination, yielding Ade+ recombinants with a strong bias in favour of deletions (Figure 1D). In contrast, Cas9d induced much lower levels of gene conversions and deletions (Figure 1D and Supplementary Table 1). Intriguingly, whilst the quantities of deDSBs and frequency of gene conversions induced by each active form of Cas9 were comparable, Cas9 induced ∼6 – 9 times more deletions than either version of Cas9n (Figure 1D and Supplementary Table 1). Altogether these data show that both lead- and lag-SSBs induced by Cas9n lead to the formation of deDSBs and elevated levels of direct repeat recombination. They also suggest that deDSBs that are induced indirectly by Cas9n are less prone to causing deletions than those induced directly by Cas9.

### DSBs arising from Cas9n-induced SSBs are primarily repaired through Rad51- dependent sister chromatid recombination

The formation of deDSBs from Cas9n-induced SSBs is most likely a consequence of replication run-off followed by fork convergence at the site of the broken fork (Supplementary Figure S1A). This process leads to one broken and one intact sister chromatid. In contrast, DSBs induced by Cas9 are independent of DNA replication and can occur simultaneously in both sister chromatids. These distinct modes of formation will have a profound impact on how the DSBs are repaired. Cas9-induced DSBs will typically be repaired by NHEJ and/or alternative end joining (a-EJ), although they can also be repaired through homology-directed pathways such as single-strand annealing (SSA)^31,32^. Indeed, the very high frequency of Cas9-induced Ade+ deletions, reported in Figure 1D, most likely arise via SSA. This mechanism occurs when DNA resection from the DSB exposes complementary ssDNA sequences at the two distinct *ade6^-^* heteroalleles, which then anneal leading to the loss of one repeat together with the DNA between the repeats^19^ (Supplementary Figure S1B). In contrast to Cas9-induced DSBs, those induced indirectly by Cas9n are more likely to undergo error-free sister chromatid exchange, which makes use of the intact sister chromatid as a template for the repair reaction. This process will occasionally give rise to Ade+ recombinants when Rad51 inadvertently catalyses pairing and strand invasion between the *ade6^-^*repeats or when resection proceeds far enough to enable SSA to occur (Supplementary Figure S1A).

To confirm our assumptions about the pathways responsible for repairing Cas9/Cas9n-induced DSBs, we first investigated whether Cas9-induced DSBs are more likely to be repaired by NHEJ/a-EJ than those induced by Cas9n. As both NHEJ and a-EJ are associated with much higher mutation rates than HR, we compared the ability of Cas9, Cas9n^H840A^, and Cas9n^D10A^ to induce mutations in a *kan^R^* gene, conferring resistance to the antibiotic geneticin (G418) (Figure 2A and 2B). Expression of Cas9, along with a gRNA that binds to the 5’ end of *kan^R^*, led to ∼90% of colonies becoming sensitive to G418 due to mutations at the Cas9 cleavage site. In contrast, neither version of Cas9n resulted in an increase in the frequency of G418 sensitive colonies (Figure 2A and 2B). These findings are consistent with the notion that most Cas9-induced DSBs are repaired by NHEJ and/or a-EJ, whereas the majority of Cas9n-induced DSBs are repaired by error-free HR.

**Figure 2.**
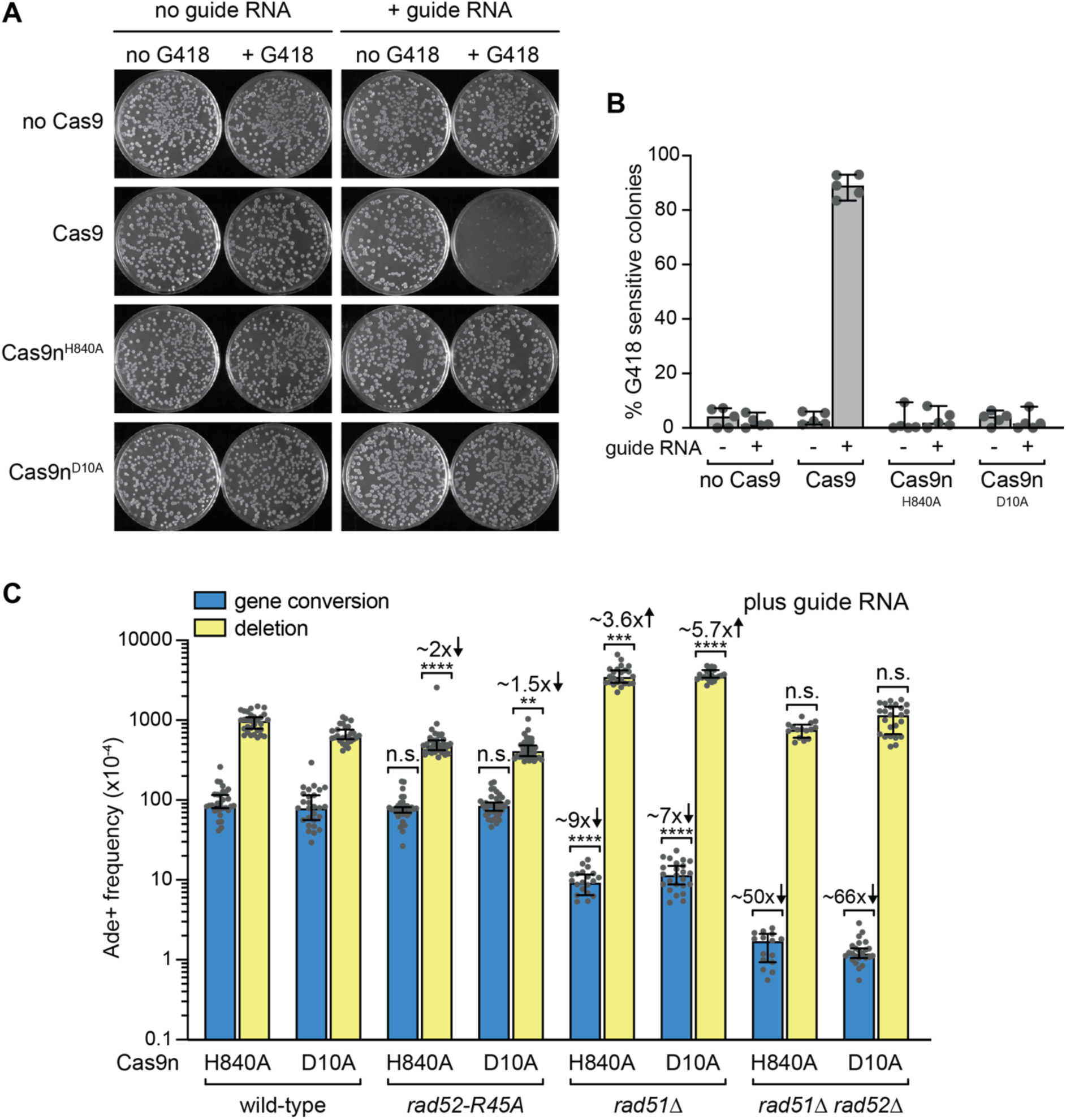
Different pathways repair the DSBs made by Cas9 and Cas9n. (**A**) Comparison of *kan^R^* gene mutation by Cas9 and Cas9n. Colonies, expressing the indicated version of Cas9, were grown with and without a gRNA that binds to the 5’ end of *kan^R^*. The colonies were then replica plated onto media with and without the G418. The images show example replica plates after incubation at 30°C for 24 hours. (**B**) Percentage of G418 sensitive colonies obtained from the plating assay shown in A. Data from 5 independent experiments are presented as mean values ± SD. Individual data points are shown as grey dots. (**C**) Frequency of Ade+ recombinants in strains containing the *ade6^-^* direct repeat substrate shown in Figure 1A. The presence of gRNA and version of Cas9n is indicated. Data are presented as median values ± 95% confidence interval with individual data points shown as grey dots. Fold changes are relative to the equivalent wild-type strain. *P*-values are for the comparison to the equivalent wild-type strain and were calculated by the Kruskal- Wallis test with Dunn’s multiple comparisons post-test. **** *p*-value <0.0001; *** *p*-value <0.001; ** *p*-value <0.01; n.s. not significant. The data are also reported in Supplementary Table 1, which includes the strain numbers, the number of colonies tested for each strain(*n*) and relevant *p*-values.

We next investigated which pathways were responsible for the high frequency of direct repeat recombination induced by Cas9n (Figure 1D). As mentioned above, deletion Ade+ recombinants can be formed by SSA, which is a pathway that is partially dependent on the ssDNA annealing activity of Rad52^33–36^. To determine whether Rad52’s ssDNA annealing activity is required for the formation of Cas9n-induced deletions, we measured the frequency of Ade+ recombinants in a *rad52*-R45A mutant that is defective for DNA annealing but proficient for Rad51-dependent recombination^37–40^. Whilst the frequency of Cas9n-induced gene conversions was unaffected by the *rad52*-R45A mutation, the frequency of deletions was reduced by up to 2-fold suggesting that some of the DSBs may indeed be repaired by SSA (Figure 2C and Supplementary Table 1).

If the majority of Cas9n-induced DSBs are repaired by Rad51-mediating entopic recombination between sister chromatids, then loss of Rad51 should result in more DSBs being repaired by SSA leading to an increase in Ade+ deletions. In agreement with this prediction, the frequency of deletions increased by up to 5.7-fold in a *rad51*Δ mutant (Figure 2C and Supplementary Table 1). These extra deletions were dependent on Rad52 suggesting that repair events that are normally mediated by Rad51 were indeed being channelled into SSA (Figure 2C and Supplementary Table 1). Notably, there were still high levels of deletions in a *rad51*Δ *rad52*Δ double mutant, which likely stem from Rad52- independent SSA. Rad51 is well-known to be the main driver of gene conversions in eukaryotes and, accordingly, we observed a ∼7 - 9-fold reduction in Cas9n-induced gene conversions in a *rad51*Δ mutant (Figure 2C and Supplementary Table 1). The gene conversions were further reduced in a *rad51*Δ *rad52*Δ double mutant suggesting that Rad52 can catalyse the formation of some gene conversions without Rad51 (Figure 2C and Supplementary Table 1). Surprisingly, even in a *rad51*Δ *rad52*Δ double mutant, we could still detect some Cas9n-induced gene conversions suggesting that there is a DSB repair pathway, which is independent of these core HR proteins, that can drive conversion or reversion of *ade6-M375*/*ade6-L469*.

Altogether these data suggest that the majority of DSBs that derive from Cas9n- induced SSBs are repaired by Rad51-dependent sister chromatid exchange. However, they also show that DSB repair can be achieved by Rad51-independent pathways, including SSA and a process that can generate gene conversions/reversions.

### Cas9n-induced SSBs can trigger BIR

Template switching is a characteristic feature of BIR, which can be detected using a direct repeat reporter positioned downstream of the DSB from where BIR is initiated^40–43^. To investigate whether any of the DSBs that derive from Cas9n-induced SSBs are repaired by BIR, we positioned the *ade6^-^* direct repeat reporter 0.7 kb downstream of a *kan^R^* gene on chromosome 3 so that we could determine the frequency of template switching through the appearance of Ade+ recombinants (Figure 3A). In the absence of gRNA, expression of Cas9 or its mutant derivatives had no effect on the frequency of Ade+ recombinants (Figure 3B). We also saw no change in the frequency of Ade+ recombinants with expression of Cas9 and Cas9d in the presence of a gRNA that binds to the *kan^R^* lagging strand template (Figure 3B). However, with Cas9n^H840A^ and Cas9n^D10A^ we observed marked increases in the frequency of both Ade+ gene conversions (5-fold and 18-fold, respectively) and Ade+ deletions (26-fold and 77-fold, respectively) suggesting that both lead- and lag-SSBs can induce BIR (Figure 3B).

**Figure 3.**
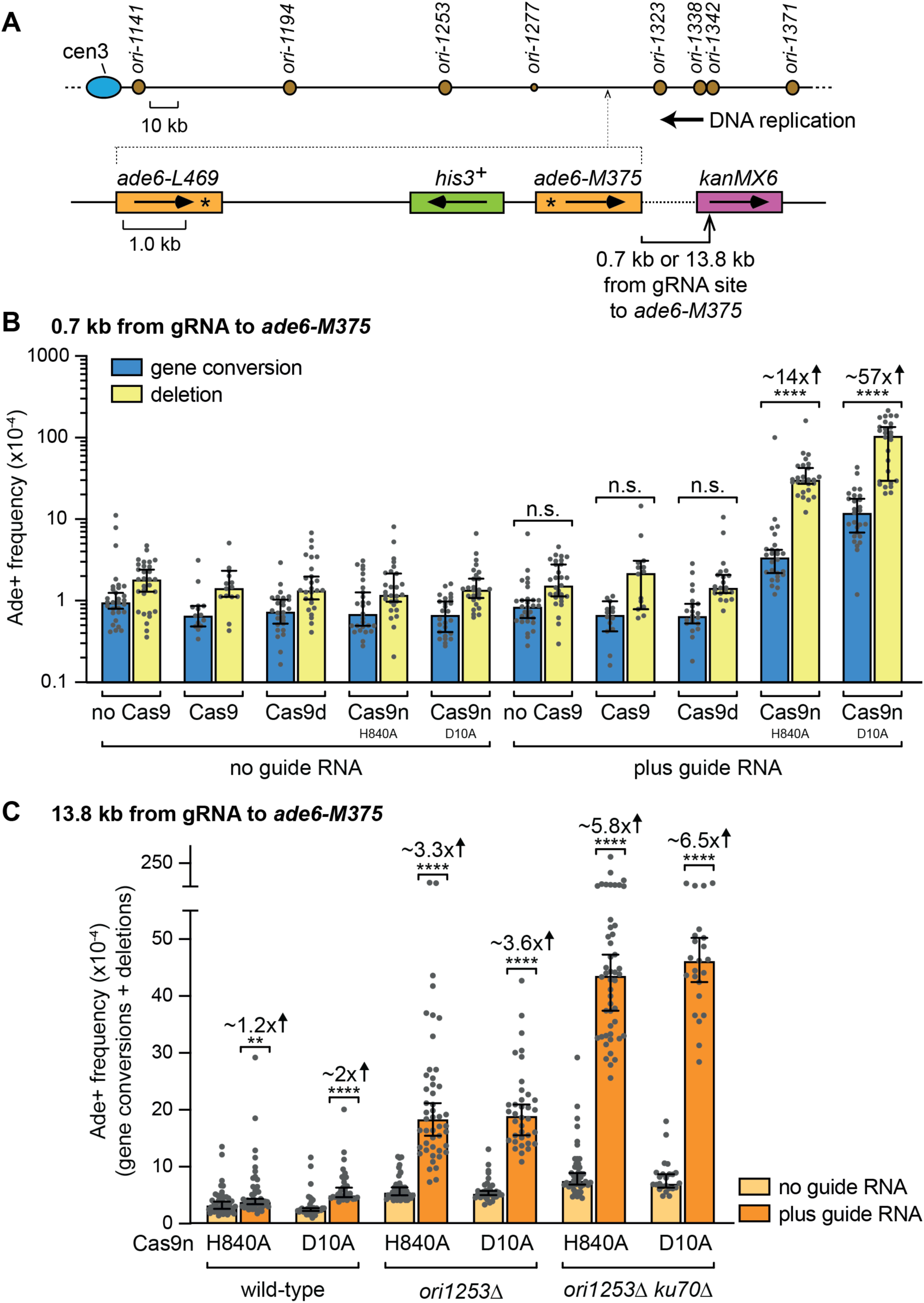
Cas9n-induced lead- and lag-SSBs trigger BIR. **(A)** Location of *kan^R^* gene (purple rectangle) upstream of the *ade6^-^* template switch reporter on chromosome 3. **(B** and **C)** Frequency of Ade+ recombinants in strains containing the *ade6^-^* template switch reporter either 0.7 kb (B) or 13.8 kb (C) downstream of the gRNA binding site in *kan^R^*. The presence of gRNA and version of Cas9 is indicated. Data are presented as median values ± 95% confidence interval with individual data points shown as grey dots. Fold changes are for the total Ade+ frequency relative to the equivalent no gRNA control strain. *P*-values are for the comparison to the equivalent no gRNA control strain and were calculated by the Mann-Whitney test (two-tailed). **** *p*-value <0.0001; ** *p*-value <0.01; n.s. not significant. The data are also reported in Supplementary Table 1, which includes the strain numbers, the number of colonies tested for each strain (*n*) and relevant *p*-values.

### BIR is limited by fork convergence

To investigate the extent of the BIR tract induced by Cas9n, we moved the *kan^R^* gene 13.8 kb upstream of the *ade6^-^* direct repeat reporter (Figure 3C). In this scenario, we observed only ∼1.2-fold increase in the frequency of Ade+ recombinants with Cas9n^H840A^ and a ∼2- fold increase with Cas9n^D10A^ (Figure 3C).

Fork convergence is known to limit BIR induced from a lead-SSB in *S. cerevisiae*^14^, to investigate whether the same holds true for lead- and lag-SSBs in *S. pombe*, we deleted the origin of replication *ori1253*, which is responsible for the majority of replication forks that converge on *kan^R^* in the centromere to telomere direction. In an *ori1253*Δ mutant, there will be more time for BIR to initiate and progress to the downstream *ade6^-^*direct repeat reporter as, in most cells, the converging replication fork now has to travel from one of the more distant centromere proximal origins (Figure 3A). In this scenario, both Cas9n variants induced a >3-fold increase in Ade+ recombinants in the presence of the gRNA indicating that BIR had had sufficient time to progress to the downstream reporter in some cells (Figure 3C). These data indicate that both lead- and lag-SSBs can give rise to BIR, which can travel at least 13.8 kb from its initiating DNA lesion if not met by an opposing replication fork.

### Ku70 is a barrier to BIR

The Ku70-Ku80 (Ku) heterodimer has been implicated in binding reversed and broken replication forks where it acts to limit DNA resection independently of its canonical role in NHEJ^44–51^. To investigate whether Ku is a barrier to Cas9n-induced BIR, we assayed template switching in a *ku70*Δ *ori1253*Δ mutant using the *ade6^-^*direct repeat reporter positioned 13.8 kb downstream of *kan^R^* (Figure 3A). In the presence of the gRNA, both versions of Cas9n induced higher levels of Ade+ recombinants in the *ku70*Δ *ori1253*Δ mutant than observed in the *ori1253*Δ single mutant (Figure 3C). These data indicate that Ku is a barrier to BIR induced by both lead- and lag-SSBs.

### Lead- and lag-SSBs induced by Flp^H305L^ and gpII trigger BIR

To investigate whether other types of SSB induce BIR, we used two different sequence-specific nicking enzymes, Flp^H305L^ and gpII from M13 bacteriophage. Flp^H305L^ is a mutant form of the Flp site-specific recombinase that nicks DNA at Flp recognition target (*FRT*) sites via a transesterification reaction that results in a SSB with Flp^H305L^ covalently linked to its 3’-phosphoryl terminus opposite a 5’-hydroxyl terminus^11,52^. These so-called Flp-nicks mimic Top1ccs that are induced by CPT. In contrast, gpII is thought to bind covalently to the 5’ termini of the nicks that it generates^10,12,53^.

We first constructed strains with *FRT* or gpII cleavage sites inserted between the *ade6^-^* heteroalleles of our direct repeat recombination reporter, with cleavage site orientation determining whether the SSB would be in the leading or lagging strand template (Figure 4A). The frequency of Ade+ recombinants in these strains was then assayed both with and without Flp^H305L^/gpII expression (Figure 4B). In all cases, expression of the nicking enzyme resulted in a strong increase in the frequency of Ade+ recombinants, ranging from a 267-fold increase for the lead-SSB induced by Flp^H305L^ to a 538-fold increase for the lag-SSB induced by gpII. These fold increases were similar to those obtained with Cas9n (Figure 4B).

**Figure 4.**
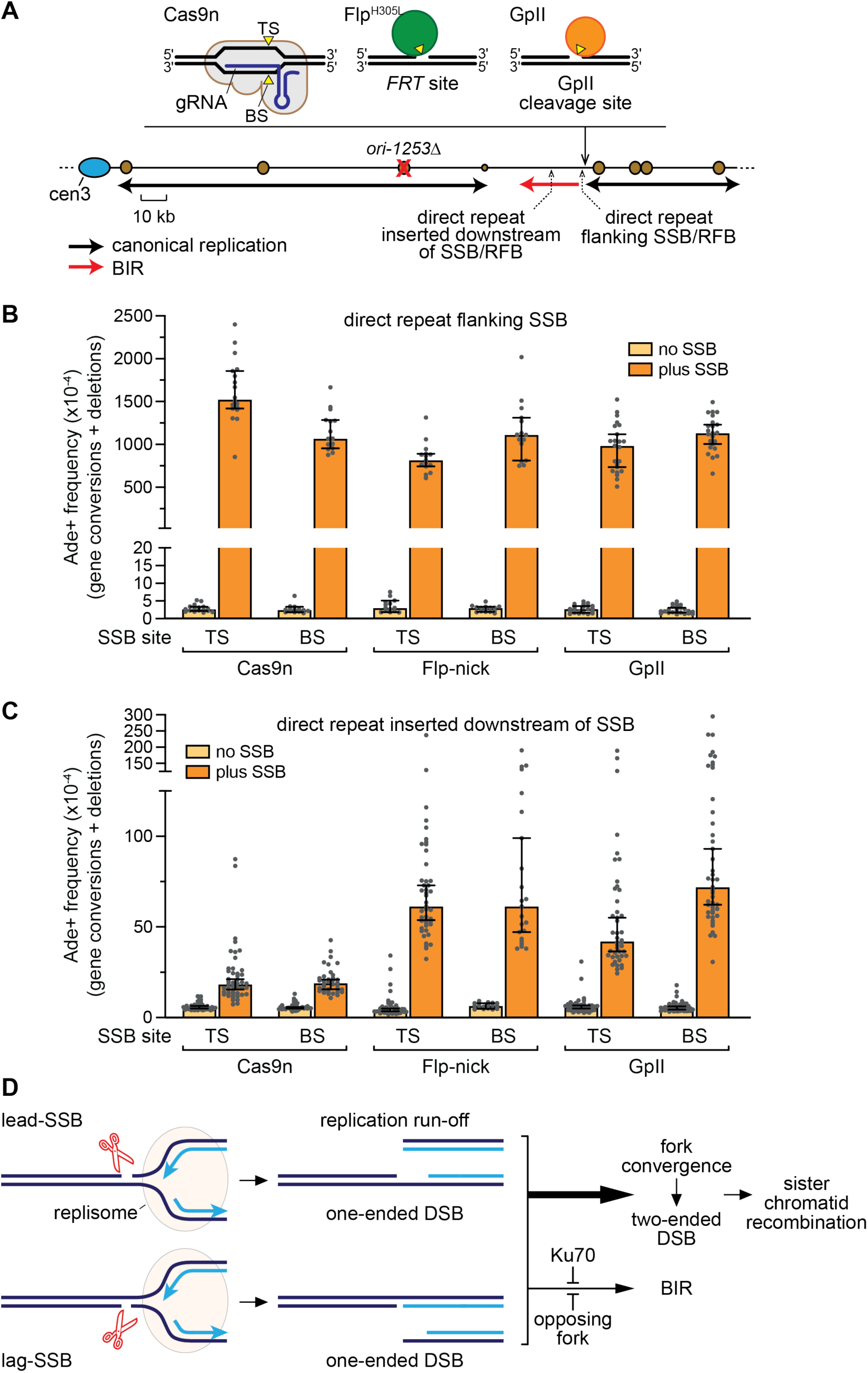
Lead- and lag-SSBs induced by Flp^H305L^ and gpII trigger BIR. **(A)** Location of the SSBs made by the three different nicking enzymes on chromosome 3. **(B** and **C)** Frequency of Ade+ recombinants in strains containing the *ade6^-^* direct repeat substrate either flanking (B) or 13.4 - 13.8 kb (C) downstream of the SSB. The position of the SSB in the leading template (top) strand (TS) and lagging template (bottom) strand (BS) is indicated. Data are presented as median values ± 95% confidence interval with individual data points shown as grey dots. The data are also reported in Supplementary Table 1, which includes the strain numbers, the number of colonies tested for each strain (*n*) and relevant *p*-values. (**D**) Model showing the different pathways that repair lead- and lag-collapsed forks.

We next compared the levels of template switching induced by Cas9n, Flp-nick, and gpII using strains with the *ade6^-^*direct repeat reporter positioned ∼13.4 – 13.8 kb downstream of the SSB and *ori1253* deleted (Figure 4A and 4C). Expression of both Flp^H305L^ and gpII resulted in a marked increase in the frequency of Ade+ recombinants ranging from a ∼7-fold increase for a lead-SSB made by gpII to a ∼15.7-fold increase for a lead-SSB made by Flp^H305L^ (Figure 4C). These increases were significantly more than the ∼3-fold increase obtained with Cas9n (Figure 4C). Altogether these data show that three different types of lead- and lag-SSBs are able to give rise to BIR suggesting that this is a universal response to fork-SSB encounters.

## Discussion

Our data indicate that encounters between replication forks and SSBs primarily give rise to deDSBs (Figure 4D). These deDSBs most likely stem from fork convergence at the SSB, resulting in the formation of one broken and one intact sister chromatid^13,54^. This is the ideal scenario for HR to deliver accurate DNA repair and, accordingly, our genetic data indicate that the majority of these deDSBs are repaired faithfully by Rad51-mediated sister chromatid exchange (Figure 4D). However, the high frequency of gene conversions and deletions between *ade6^-^*heteroalleles that flank the deDSB demonstrate that HR is not always accurate and can inadvertently recombine repetitive DNA sequences as has been shown in previous studies. Interestingly, we find that Rad52’s ssDNA annealing activity is required for some of the deletions that are formed when Rad51 is present. Its role here might be to promote the formation of a double Holliday junction by mediating second end capture^55–58^. Alternatively, these deletions may stem from Rad52-mediated SSA, which is known to compete with Rad51-dependent DSB repair^34–36^.

Even though the majority of fork-SSB encounters appear to be resolved through fork convergence and deDSB repair, our data indicate that some lead to BIR. Importantly, we show for the first time that both lead- and lag-SSBs are capable of triggering BIR. This discovery is consistent with the recent finding that fork encounters with both lead- and lag- SSBs give rise to seDSBs^15^. The termini of these seDSBs differ depending on whether they stem from fork collapse at a lead- or lag-SSB. Lead-collapse generates almost blunt seDSBs, whereas lag-collapse generates a seDSB with a ∼70 nucleotide 3’ ssDNA overhang^15^. Moreover, lead-collapse generates an unbroken sister chromatid with a ssDNA gap between the final primed Okazaki fragment and the 3’ terminus of the SSB, whereas lag-collapse generates an unbroken sister chromatid with just a nick^15^. It has been suggested that these differences could have an impact on the subsequent repair. Repair of lead- collapse seDSBs might be more reliant on gap filling and DNA end resection, whereas lag- collapse seDSBs would require no gap filling and have a ready-made 3’ ssDNA tail sufficient for Rad51-mediated strand invasion^15^. Such differences could also have an impact on the speed at which BIR is initiated, with the additional processing requirements of lead-collapsed forks resulting in slower BIR initiation. This could explain why we observe a higher frequency of template switching when our recombination reporter is positioned 0.7 kb downstream of a lag-SSB compared to a lead-SSB (Figure 3B). However, this difference appears to dissipate when the reporter is positioned 13.8 kb downstream of the SSB and the oncoming replication fork is delayed by deleting *ori1253* (Figure 3C). It has also been suggested that Ku might have a bigger influence on the processing of lead- collapsed forks than those that collapse at lag-SSBs because of its preference for binding blunt or nearly blunt dsDNA ends^15^. However, we observe an almost identical increase in BIR-associated templating switching with both lead- and lag-SSBs when *ku70* is deleted (Figure 3C). This suggests that Ku is a barrier to BIR regardless of whether it is initiated from a lead- or lag-collapse. It should be noted that, whilst Ku binds less efficiently to ssDNA than dsDNA, it can localize to ssDNA-dsDNA junctions and protect 5’-recessed ends from degradation by Exo1 in vitro^59^.

When measured in budding yeast outside of S-phase, BIR is found to be quite a slow process with DNA synthesis of the first 500 bp detected 2.5 hours after DSB induction^60^. Once underway, the average rate of DNA synthesis by BIR is 0.5 kb per minute, which is four-fold slower than the average velocity of a canonical replication fork in *S. cerevisiae*^60,61^. The slow kinetics of BIR is likely to be a key limiting factor for its deployment in repairing broken replication forks, as in most cases fork convergence will occur before BIR has begun. In our experimental system, template switching 13.8 kb downstream of the SSB was only detected when fork convergence was delayed by deleting *ori1253*. In this scenario, the converging fork originates predominantly from either *ori1194* or *ori1141* (Figure 3A). Assuming that at least one of these origins fire during early S-phase, and the resulting fork moves with an average velocity of 2 kb per minute, then BIR would have less than 1.5 hours to register a template switch at the downstream reporter^42^. Either BIR has faster kinetics when induced from a broken replication fork in *S. pombe* than from a non-S-phase DSB in *S. cerevisiae*, or the template switches that we detect only occur in cells where the converging fork undergoes further delay.

SSBs made by Flp-nick and gpII induce more template switching than those made by Cas9n (Figure 4C). Considering that Cas9n, Flp-nick and gpII induce similar levels of direct repeat recombination when their target cleavage site is flanked by the *ade6^-^* heteroalleles (Figure 4B), the higher levels of template switching observed at the downstream reporter with Flp-nick and gpII compared to Cas9n suggest that some types of SSB may more readily induce BIR than others. In future studies it will be important to elucidate all of the factors that govern the deployment of BIR to ensure that this mutagenic mode of DNA synthesis is only unleashed as a pathway of last resort.

## Materials and Methods

### Yeast strains and plasmid construction

*S. pombe* strains are listed in Supplementary Table S2. Derivatives of recombination reporter strains carrying the indicated gene/replication origin deletion(s) were obtained from genetic crosses. Strain MCW9804 was constructed by transformation of BlpI- linearized plasmid pLX27 into a *ade6-M375* strain^62^. pLX27 is a derivative of pCB33 in which the *RTS1* sequence has been removed by SacI digestion^63^. Plasmids pLX1, pLX5, pLX6 and pLX10 are derivatives of pAG32^64^, which were used for the targeted integration of P*nmt81*-*cas9*/*cas9n*/*cas9d*-*hphMX4* at the *lys1* locus. pLX1 contains *cas9n^D10A^*, which was cloned from pSPCas9n(BB)-2A-GFP (Addgene plasmid ID: #48140)^65^. pLX5 (*cas9d^D10A/H840A^*), pLX6 (*cas9n^H840A^*), and pLX10 (*cas9*), were derived from pLX1 by site-directed mutagenesis of *cas9n^D10A^* to obtain the required version of Cas9. All versions of Cas9 were expressed from the thiamine repressible *nmt81* promoter^66^. Plasmid pLX2 is a derivative of pMZ374 (Addgene plasmid ID: #59896)^67^ in which the Cas9 gene has been removed by EagI digestion, and annealed oligonucleotides, oMW1708 (5’- AAGCTTATCGATACCGTCGAGT-3’) plus oMW1709 (5’-TCGACGGTATCGATAAGCTTTT-3’), inserted at the CspCI site. This plasmid contains a *ura4^+^* marker and expresses a gRNA that anneals to the lagging strand template in the region adjacent to the 5’ end of the *his3* gene in the *ade6^-^* direct repeat recombination reporter shown in Figure 1A. pLX3 is a derivative of pLX2 that lacks the sequence for the gRNA. It was constructed by digesting pLX2 with BamHI and NheI to remove the gRNA sequence, and then re-ligating the DNA ends after blunt ending them with DNA Polymerase I, Large (Klenow) Fragment (New England BioLabs, M0210L). pLX32 is a derivative of pMZ283 (Addgene plasmid ID: #52224)^67^, which was obtained by site-directed mutagenesis of pMZ283 using primers oMW2158 (5’- GTCTTTTCCTTTCTTCGGTACAGGTTATG-3’) and oMW2159 (5’-TCACGTTTCGGTTTTAGAGCTAGAAATAGC-3’). This plasmid contains a *ura4^+^* marker and expresses a gRNA that anneals to the lagging strand template at the 5’ end of the *kan^R^* gene shown in Figure 3A. pLX46 and pLX47 are derivatives of pLX32 and pMZ283, respectively, which contain a *LEU2* gene instead of *ura4*. The construction of the *ade6^-^* direct repeat used to monitor template switching has been described previously ^40,43^. Placement of the *kan^R^* gene at the *ade6* locus, ∼13.8 kb upstream of this reporter, was done by targeted gene replacement using a NotI restriction fragment from pLX44. pLX44 was derived from pMW921^42^ by swapping a SalI-SacI fragment containing *RTS1*-*hphMX4* for the *kanMX6* cassette from pFA6aKanMX6^68^. Placement of *FRT* sites and gpII cleavage sites upstream of the template switch reporter was also done by targeted gene replacement of *ade6* using NotI restriction fragments from plasmids pYL8 (gpII TS cleavage site), pYL9 (gpII BS cleavage site), pKT1 (*FRT* BS cleavage site), and pKT2 (*FRT* TS cleavage site). These plasmids were made by swapping a SalI-SmaI fragment in pMW921, containing *RTS1*, for the following annealed oligonucleotides: oMW480 (5’- TTTAAAGTCGACCTCGAGGTTCTTTAATAGTGGACTCTTGTTCCAAACTGGAACAACACCCGGGTTTAAA-3’) and oMW481 (5’- TTTAAACCCGGGTGTTGTTCCAGTTTGGAACAAGAGTCCACTATTAAAGAACCTCGAGGTCGACTTTAAA-3’) (pYL8); oMW482 (5’- TTTAAACCCGGGCTCGAGGTTCTTTAATAGTGGACTCTTGTTCCAAACTGGAACAACAGTCGACTTTAAA-3’) and oMW483 (5’-TTTAAAGTCGACTGTTGTTCCAGTTTGGAACAAGAGTCCACTATTAAAGAACCTCGAGCCCGGGTTTAAA-3’) (pYL9); oMW2031 (5’- ATCGAAGTTCCTATACTTTCTAGAGAATAGGAACTTCCGAATAGGAACTTCG- 3’) and oMW2032 (5’- TCGACGAAGTTCCTATTCGGAAGTTCCTATTCTCTAGAAAGTATAGGAACTTC GAT-3’) (pKT1); oMW2068 (5’- ATCGAAGTTCCTATTCGGAAGTTCCTATTCTCTAGAAAGTATAGGAACTTCG- 3’) and oMW2069 (5’- TCGACGAAGTTCCTATACTTTCTAGAGAATAGGAACTTCCGAATAGGAACTTC GAT-3’) (pKT2). Strains MCW10107 and MCW10108, which contain the *ade6^-^*direct repeat recombination reporter with an *FRT* site between *his3* and *ade6-M375*, were made by transformation of BlpI-linearised plasmid pYL7 and pYL4, respectively, into a *ade6- M375* strain. Both plasmids are derivatives of pFOX2^62^ with annealed oligonucleotides, oMW2031 plus oMW2032 (pYL7) or oMW2068 plus oMW2069 (pYL4), inserted between SalI and SmaI restriction sites. The plasmids for expression of Flp^H305L^ and gpII are derivatives of pREP1 ^69,70^. pREP1- Flp^H305L^ (pPC7) was made by first amplifying the Flp^H305L^ gene from plasmid pFV17D-H305L^11^ using primers oMW2029 (5’- GGGTCGACATGCCACAATTTGGTATATTATGTA-3’) plus oMW2030 (5’- TAGGATCCTAATTATATGCGTCTATTTATGTAGGATGAAA-3’), and then cloning it as a SalI-BamHI fragment into pREP1 so that its expression would be directed by the thiamine repressible *nmt1* promoter. The construction of pREP1-NLS-gpII has been described^12^. Plasmids were verified by DNA sequencing. The genotype of each *S. pombe* strain was confirmed by determining the presence/absence of relevant genetic markers, as well as diagnostic PCRs and DNA sequencing where appropriate.

### Media and growth conditions

Standard protocols were used for the growth and genetic manipulation of *S. pombe*^71^. The complete and minimal media were yeast extract with supplements (YES) and Edinburgh minimal medium plus 3.7 g/l sodium glutamate (EMMG) and appropriate amino acids (225 mg/l), respectively. Strains carrying the integrated P*nmt81*-*cas9/cas9n/cas9d* constructs or pREP1/pREP1-FlpH305L/gpII were grown on EMMG supplemented with thiamine (4 µM final concentration) to repress expression from the *nmt* promoter. Strains carrying plasmids were grown on EMMG lacking leucine or uracil as appropriate. To maintain the integrity of the *ade6-L469* – *ade6-M375* recombination reporters, strains carrying them were grown on EMMG lacking histidine and supplemented with low levels of adenine (10 mg/l) to distinguish non-recombinant colonies (red) from Ade+ recombinants (white). For recombination assays, total number of colony forming cells was determined by plating onto YES supplemented with low levels of adenine (10 mg/l) (YES/LA). Ade+ recombinants were selected on YES lacking adenine and supplemented with 200 mg/l of guanine (YES- ade+gua) to prevent uptake of residual adenine^72^. All strains were grown at 30°C.

### G418 sensitivity assay

Relevant strains carrying the *kan^R^* gene adjacent to the *ade6^-^* direct repeat recombination reporter on chromosome 3 (Figure 3A), were transformed with the gRNA expressing plasmid pLX32 or the no gRNA control plasmid pLX3. Five independent transformants for each strain were plated onto EMMG agar lacking uracil and thiamine. Colonies were grown for 6 days at 30°C before being replica plated onto YES and YES containing G418 (100 mg/l). Plates were incubated for 24 hours at 30°C before being photographed

### Recombination assays

The protocol for determining the frequency of Ade^+^ recombinants has been described^72^. Strains transformed with the appropriate gRNA/Flp^H305L^/gpII/control plasmid were grown for 6 - 10 days on EMMG plates containing either uracil or leucine for plasmid selection, but lacking thiamine to allow for Cas9/Cas9n/Cas9d/Flp^H305L^/gpII expression and the induction of recombination. Colonies from these plates were then analysed for their frequency of Ade^+^ recombinant cells. This was done by serially diluting each colony in sterile Milli-Q water and then plating appropriate dilutions on YES/LA and YES-ade+gua. These plates were grown for 5 - 6 days. Colonies were then counted using a Protos 3 automated colony counter (Synbiosis) with Protos 3 Version 1.2.4.0 software (Synoptics Ltd). The frequency of Ade+ recombinants amongst total viable (colony forming) cells was determined by comparing the number of colonies on the YES/LA plate with those on the YES-ade+gua plate. The percentage of deletions and gene conversions amongst Ade+ recombinants were determined by replica plating the YES-ade+gua plates onto EMMG plates lacking histidine. For each strain, at least three independent gRNA/Flp^H305L^/gpII/control plasmid transformants were assayed and, for each of these transformants, the recombination frequency was measured in 4 - 15 colonies. A single recombination event at an early stage in the growth of a colony will give rise to many recombinant cells. To avoid these so-called jackpot events from skewing the data, we used the method of the median to determine the recombination frequency for each strain.

### Statistical analysis

Analysis of recombination data was performed using Excel for Mac Version 16.78.3 (Microsoft®) GraphPad Prism Version 10.1.0 (GraphPad Software, San Diego, CA). Recombination frequencies were analysed for normal distribution using the Shapiro-Wilk test. Not all data passed this test and, therefore, the data were evaluated for lognormal distribution using the Shapiro-Wilk test. Data that conformed to a lognormal distribution were log-transformed and relevant recombination frequencies were compared using an Unpaired t-test (two-tailed). For data that did not follow a lognormal distribution, recombination frequencies were compared using a Two-tailed Mann Whitney test or a Kruskal-Wallis test (one-way ANOVA on ranks) with a Dunn’s multiple comparisons post-test. These are non-parametric statistical tests and, therefore, do not require the data to be normally distributed. *p*-values are reported in Supplementary Table S1 and Supplementary Data file 1.

### DSB detection

Strains were grown in shake flasks containing 500 ml EMMG (lacking uracil and thiamine) for 48 - 50 hours reaching a density of ∼1x10^7^ cells/ml. The cultures were then treated with 0.1% sodium azide and harvested by centrifugation. Cell pellets were washed in 20 ml 50 mM EDTA and stored in two equal sized aliquots at -80°C to await further processing. Cells from one aliquot were defrosted on ice and gently resuspended in an equal volume of CPES buffer to which *Trichoderma harzianum* lysing enzymes (250 mg/ml; Sigma-Aldrich, L1412), *Arthrobacter luteus* lyticase (25 mg/ml; Sigma-Aldrich, L4025), Zymolyase 20T (100 mg/ml; MP Biomedicals, 08320922), and DTT (10 µl/ml of a 1 M solution; Sigma-Aldrich, D9779) were added. The suspension was then incubated at 37°C for up to 2 hours until spheroplasting of the cells was achieved. The spheroplasted cells were then mixed with an equal volume of molten low-melting point agarose (2%; Thermo Scientific, R0801) in 0.25 M EDTA and 1.2 M sorbitol. The cell/agarose mixture was pipetted into ∼30 plug molds (BioRad, 1703713) and allowed to set at 4°C for 15 minutes. The plugs were then removed from the molds and incubated for 36 hours at 50°C in 3 ml of NDS-PK buffer (10 mM Tris-HCl pH 7.5, 1% N-lauroylsarcosine, 495 mM EDTA, 0.5 mg/ml Proteinase K), with the buffer being replaced with a fresh 3 ml aliquot after 18 hours. Following removal of the NDS-PK buffer, the plugs were washed for 1 hour in 10 ml of 1x TE buffer (10 mM Tris-HCl pH 8.0, 1 mM EDTA) containing 0.5 mM PMSF, followed by three sequential 1-hour wash steps in 20 ml of 1x TE. The plugs were then kept overnight at 4°C in 1x TE. For all the following steps, DNA LoBind tubes (Eppendorf: 1130122232; 0030122208; 022431021) were used. The plugs were washed in 20 ml of Milli-Q water for 30 minutes and then 10 ml of 1x rCutSmart buffer (New England Biolabs, B6004S) for 1 hour followed by a further hour in 5 ml of 1x rCutSmart buffer. The buffer was then removed and the plugs were melted by placing the tube at 65°C in a water bath for 15 - 20 minutes. The tube containing the plugs was then cooled to 42°C before adding 5 µl of Beta-Agarase I (New England Biolabs, M0392) and incubating at 42°C for 1 hour. 200 units of PstI-HF (New England Biolabs, R3140L) plus 200 units of NheI-HF (New England Biolabs, R3131L), and 20 µl of RNase A (100 mg/ml; Qiagen; 1007885) were then added and the mixture was incubated overnight at 37°C. An extra 100 units of PstI-HF and NheI-HF were added the next day and the incubation continued for a further 6 hours. The mixture was then centrifuged at 3,200 x g for 10 minutes and the supernatant transferred to a fresh tube. The supernatant was then mixed with an equal volume of phenol:chloroform:isoamyl alcohol (25:24:1 v/v; ThermoFisher Scientific, 15593031), and centrifuged for 10 minutes. DNA in the aqueous phase was then precipitated by addition of an equal volume of isopropanol plus 0.1 volume of 3 M sodium acetate and incubation overnight at -20°C. Following centrifugation, the DNA pellet was washed three times in 70% ethanol, dried and dissolved in 40 µl of 1x TE buffer. The sample was then mixed with 4 µl of 10 x gel loading dye (50% glycerol, 1% SDS, 0.1 M EDTA, 0.25% bromophenol blue), and equal sized aliquots (∼10 µg of DNA in each) were then run on separate 1% agarose gels in 1x TBE buffer for 24 hours at 50 V. The gel was then stained with Ethidium Bromide for 1 hour to visualise the DNA before Southern blotting. The gel was treated with 0.25 M HCl to depurinate the DNA, and then transferred to GeneScreen Plus membrane (Perkin Elmer, NEF988001PK) by capillary action under alkaline conditions. Following transfer, the membrane was probed with either ^32^P-radiolabelled probe A or probe S (Rediprime II Random Prime Labelling System, Cytiva, RPN1633; and alpha-^32^P dCTP, Perkin-Elmer, NEG513H250UC) in ULTRA-hyb buffer (Invitrogen, AM8669) at 42°C. The membrane was then washed following the manufacturer’s recommended procedure and exposed to a phosphor screen for up to 5 days. The phosphor screen was scanned using a FLA-3000 Phosphorimager (Fujifilm) controlled by Image Reader software (Fujifilm, Version 2.02), and the data analysed using Image Gauge software (Fujifilm, Version 4.21).

## Acknowledgements

We thank Lotte Bjergbæk for the gift of plasmids. This work was supported by grants from the Medical Research Council (MR/V009214/1 awarded to M.C.W.), and Biotechnology and Biological Sciences Research Council (BB/P019706/1 and BB/V00073X/1 awarded to M.C.W.). The funders had no role in study design, data collection and analysis, the decision to publish, or preparation of the manuscript.

## Author contributions

Y.X., Y.L., C.A.M., K.T., and S.J. performed the experiments; P.C. and C.T. performed preliminary experiments; Y.X., Y.L., C.A.M., K.T., and M.C.W. contributed to experimental design, data analysis and presentation; C.A. provided expertise and feedback; M.C.W. secured funding and wrote the manuscript.

## References

1. Caldecott, K.W. (2008). Single-strand break repair and genetic disease. Nat. Rev. Genet. 9, 619–631.

2. Caldecott, K.W. (2022). DNA single-strand break repair and human genetic disease. Trends Cell Biol. 32, 733–745.

3. Pommier, Y. (2006). Topoisomerase I inhibitors: camptothecins and beyond. Nat. Rev. Cancer 6, 789–802.

4. Thomas, A., and Pommier, Y. (2019). Targeting Topoisomerase I in the Era of Precision Medicine. Clin. Cancer Res. 25, 6581–6589.

5. Hengel, S.R., Spies, M.A., and Spies, M. (2017). Small-Molecule Inhibitors Targeting DNA Repair and DNA Repair Deficiency in Research and Cancer Therapy. Cell Chem. Biol. 24, 1101–1119.

6. Avemann, K., Knippers, R., Koller, T., and Sogo, J.M. (1988). Camptothecin, a specific inhibitor of type I DNA topoisomerase, induces DNA breakage at replication forks. Mol. Cell Biol. 8, 3026–3034.

7. Ryan, A.J., Squires, S., Strutt, H.L., and Johnson, R.T. (1991). Camptothecin cytotoxicity in mammalian cells is associated with the induction of persistent double strand breaks in replicating DNA. Nucleic Acids Res. 19, 3295–3300.

8. Tsao, Y.P., Russo, A., Nyamuswa, G., Silber, R., and Liu, L.F. (1993). Interaction between replication forks and topoisomerase I-DNA cleavable complexes: studies in a cell-free SV40 DNA replication system. Cancer Res. 53, 5908–5914.

9. Strumberg, D., Pilon, A.A., Smith, M., Hickey, R., Malkas, L., and Pommier, Y. (2000). Conversion of topoisomerase I cleavage complexes on the leading strand of ribosomal DNA into 5’-phosphorylated DNA double-strand breaks by replication runoff. Mol. Cell Biol. 20, 3977–3987.

10. Kuzminov, A. (2001). Single-strand interruptions in replicating chromosomes cause double-strand breaks. Proc. Natl. Acad. Sci. U S A 98, 8241–8246.

11. Nielsen, I., Bentsen, I.B., Lisby, M., Hansen, S., Mundbjerg, K., Andersen, A.H., and Bjergbaek, L. (2009). A Flp-nick system to study repair of a single protein-bound nick in vivo. Nat. Methods 6, 753–757.

12. Osman, F., Ahn, J.S., Lorenz, A., and Whitby, M.C. (2016). The RecQ DNA helicase Rqh1 constrains Exonuclease 1-dependent recombination at stalled replication forks. Sci. Rep. 6, 22837.

13. Cortes-Ledesma, F., and Aguilera, A. (2006). Double-strand breaks arising by replication through a nick are repaired by cohesin-dependent sister-chromatid exchange. EMBO Rep. 7, 919–926.

14. Mayle, R., Campbell, I.M., Beck, C.R., Yu, Y., Wilson, M., Shaw, C.A., Bjergbaek, L., Lupski, J.R., and Ira, G. (2015). Mus81 and converging forks limit the mutagenicity of replication fork breakage. Science 349, 742–747.

15. Vrtis, K.B., Dewar, J.M., Chistol, G., Wu, R.A., Graham, T.G.W., and Walter, J.C. (2021). Single-strand DNA breaks cause replisome disassembly. Mol. Cell 81, 1309–1318.

16. Gonzalez-Barrera, S., Cortes-Ledesma, F., Wellinger, R.E., and Aguilera, A. (2003). Equal sister chromatid exchange is a major mechanism of double-strand break repair in yeast. Mol. Cell. 11, 1661–1671.

17. Cortes-Ledesma, F., Tous, C., and Aguilera, A. (2007). Different genetic requirements for repair of replication-born double-strand breaks by sister-chromatid recombination and break-induced replication. Nucleic Acids Res. 35, 6560–6570.

18. Mehta, A., and Haber, J.E. (2014). Sources of DNA double-strand breaks and models of recombinational DNA repair. Cold Spring Harb. Perspect. Biol. 6, a016428.

19. Al-Zain, A.M., and Symington, L.S. (2021). The dark side of homology-directed repair. DNA Repair (Amst) 106, 103181.

20. Scully, R., Panday, A., Elango, R., and Willis, N.A. (2019). DNA double-strand break repair-pathway choice in somatic mammalian cells. Nat. Rev. Mol. Cell Biol. 20, 698–714.

21. Kockler, Z.W., Osia, B., Lee, R., Musmaker, K., and Malkova, A. (2021). Repair of DNA Breaks by Break-Induced Replication. Annu. Rev. Biochem. 90, 165–191.

22. Liu, L., and Malkova, A. (2022). Break-induced replication: unraveling each step. Trends Genet. 38, 752–765.

23. Carvalho, C.M., and Lupski, J.R. (2016). Mechanisms underlying structural variant formation in genomic disorders. Nat. Rev. Genet. 17, 224–238.

24. Li, Y., Roberts, N.D., Wala, J.A., Shapira, O., Schumacher, S.E., Kumar, K., Khurana, E., Waszak, S., Korbel, J.O., Haber, J.E., et al. (2020). Patterns of somatic structural variation in human cancer genomes. Nature 578, 112–121.

25. Costantino, L., Sotiriou, S.K., Rantala, J.K., Magin, S., Mladenov, E., Helleday, T., Haber, J.E., Iliakis, G., Kallioniemi, O.P., and Halazonetis, T.D. (2014). Break-induced replication repair of damaged forks induces genomic duplications in human cells. Science 343, 88–91.

26. Sotiriou, S.K., Kamileri, I., Lugli, N., Evangelou, K., Da-Re, C., Huber, F., Padayachy, L., Tardy, S., Nicati, N.L., Barriot, S., et al. (2016). Mammalian RAD52 Functions in Break-Induced Replication Repair of Collapsed DNA Replication Forks. Mol. Cell 64, 1127–1134.

27. Minocherhomji, S., Ying, S., Bjerregaard, V.A., Bursomanno, S., Aleliunaite, A., Wu, W., Mankouri, H.W., Shen, H., Liu, Y., and Hickson, I.D. (2015). Replication stress activates DNA repair synthesis in mitosis. Nature 528, 286–290.

28. Bhowmick, R., Minocherhomji, S., and Hickson, I.D. (2016). RAD52 Facilitates Mitotic DNA Synthesis Following Replication Stress. Mol. Cell 64, 1117–1126.

29. Bhowmick, R., Hickson, I.D., and Liu, Y. (2023). Completing genome replication outside of S phase. Mol. Cell. 10.1016/j.molcel.2023.08.023.

30. Sipiczki, M. (2000). Where does fission yeast sit on the tree of life? Genome biology 1, Reviews1011. 10.1186/gb-2000-1-2-reviews1011.

31. Xue, C., and Greene, E.C. (2021). DNA Repair Pathway Choices in CRISPR-Cas9- Mediated Genome Editing. Trends Genet. 37, 639–656.

32. van de Kooij, B., Kruswick, A., van Attikum, H., and Yaffe, M.B. (2022). Multi- pathway DNA-repair reporters reveal competition between end-joining, single-strand annealing and homologous recombination at Cas9-induced DNA double-strand breaks. Nat. Commun. 13, 5295.

33. Bhargava, R., Onyango, D.O., and Stark, J.M. (2016). Regulation of Single-Strand Annealing and its Role in Genome Maintenance. Trends Genet. 32, 566–575.

34. Fishman-Lobell, J., Rudin, N., and Haber, J.E. (1992). Two alternative pathways of double-strand break repair that are kinetically separable and independently modulated. Mol. Cell Biol. 12, 1292–1303.

35. Sugawara, N., and Haber, J.E. (1992). Characterization of double-strand break-induced recombination: homology requirements and single-stranded DNA formation. Mol. Cell Biol. 12, 563–575.

36. Ivanov, E.L., Sugawara, N., Fishman-Lobell, J., and Haber, J.E. (1996). Genetic requirements for the single-strand annealing pathway of double- strand break repair in *Saccharomyces cerevisiae*. Genetics 142, 693–704.

37. Yan, Z., Delannoy, M., Ling, C., Daee, D., Osman, F., Muniandy, P.A., Shen, X., Oostra, A.B., Du, H., Steltenpool, J., et al. (2010). A histone-fold complex and FANCM form a conserved DNA-remodeling complex to maintain genome stability. Mol. Cell 37, 865–878.

38. Bai, Y., Davis, A.P., and Symington, L.S. (1999). A novel allele of RAD52 that causes severe DNA repair and recombination deficiencies only in the absence of RAD51 or RAD59. Genetics 153, 1117–1130.

39. Shi, I., Hallwyl, S.C., Seong, C., Mortensen, U., Rothstein, R., and Sung, P. (2009). Role of the Rad52 amino-terminal DNA binding activity in DNA strand capture in homologous recombination. J. Biol. Chem. 284, 33275–33284.

40. Kishkevich, A., Tamang, S., Nguyen, M.O., Oehler, J., Bulmaga, E., Andreadis, C., Morrow, C.A., Jalan, M., Osman, F., and Whitby, M.C. (2022). Rad52’s DNA annealing activity drives template switching associated with restarted DNA replication. Nat. Commun. 13, 7293.

41. Smith, C.E., Llorente, B., and Symington, L.S. (2007). Template switching during break-induced replication. Nature 447, 102–105.

42. Nguyen, M.O., Jalan, M., Morrow, C.A., Osman, F., and Whitby, M.C. (2015). Recombination occurs within minutes of replication blockage by *RTS1* producing restarted forks that are prone to collapse. Elife 4, e04539.

43. Jalan, M., Oehler, J., Morrow, C.A., Osman, F., and Whitby, M.C. (2019). Factors affecting template switch recombination associated with restarted DNA replication. Elife 8. 10.7554/eLife.41697.

44. Teixeira-Silva, A., Ait Saada, A., Hardy, J., Iraqui, I., Nocente, M.C., Freon, K., and Lambert, S.A.E. (2017). The end-joining factor Ku acts in the end-resection of double strand break-free arrested replication forks. Nat. Commun. 8, 1982.

45. Audoynaud, C., Schirmeisen, K., Ait Saada, A., Gesnik, A., Fernandez-Varela, P., Boucherit, V., Ropars, V., Chaudhuri, A., Freon, K., Charbonnier, J.B., and Lambert, S.A.E. (2023). RNA:DNA hybrids from Okazaki fragments contribute to establish the Ku-mediated barrier to replication-fork degradation. Mol. Cell 83, 1061–1074.

46. Whelan, D.R., Lee, W.T.C., Marks, F., Kong, Y.T., Yin, Y., and Rothenberg, E. (2020). Super-resolution visualization of distinct stalled and broken replication fork structures. PLoS Genet. 16, e1009256.

47. Jones, C.E., and Forsburg, S.L. (2021). Monitoring *Schizosaccharomyces pombe* genome stress by visualizing end-binding protein Ku. Biol. Open 10. 10.1242/bio.054346.

48. Chanut, P., Britton, S., Coates, J., Jackson, S.P., and Calsou, P. (2016). Coordinated nuclease activities counteract Ku at single-ended DNA double-strand breaks. Nat. Commun. 7, 12889.

49. Balestrini, A., Ristic, D., Dionne, I., Liu, X.Z., Wyman, C., Wellinger, R.J., and Petrini, J.H. (2013). The Ku heterodimer and the metabolism of single-ended DNA double-strand breaks. Cell Rep. 3, 2033–2045.

50. Foster, T.J., Lundblad, V., Hanley-Way, S., Halling, S.M., and Kleckner, N. (1981). Three Tn10-associated excision events: relationship to transposition and role of direct and inverted repeats. Cell 23, 215–227.

51. Mimitou, E.P., and Symington, L.S. (2010). Ku prevents Exo1 and Sgs1-dependent resection of DNA ends in the absence of a functional MRX complex or Sae2. EMBO J. 29, 3358–3369.

52. Nielsen, I., Andersen, A.H., and Bjergbaek, L. (2012). Studying repair of a single protein-bound nick in vivo using the Flp-nick system. Methods Mol. Biol. 920, 393–415.

53. Asano, S., Higashitani, A., and Horiuchi, K. (1999). Filamentous phage replication initiator protein gpII forms a covalent complex with the 5’ end of the nick it introduced. Nucleic Acids Res. 27, 1882–1889.

54. Kuzminov, A. (2001). DNA replication meets genetic exchange: chromosomal damage and its repair by homologous recombination. Proc. Natl. Acad. Sci. U S A 98, 8461–8468.

55. Lao, J.P., Oh, S.D., Shinohara, M., Shinohara, A., and Hunter, N. (2008). Rad52 promotes postinvasion steps of meiotic double-strand-break repair. Mol. Cell 29, 517–524.

56. McIlwraith, M.J., and West, S.C. (2008). DNA repair synthesis facilitates RAD52- mediated second-end capture during DSB repair. Mol. Cell 29, 510–516.

57. Nimonkar, A.V., Sica, R.A., and Kowalczykowski, S.C. (2009). Rad52 promotes second-end DNA capture in double-stranded break repair to form complement- stabilized joint molecules. Proc. Natl. Acad. Sci. U S A 106, 3077–3082.

58. Sugiyama, T., Kantake, N., Wu, Y., and Kowalczykowski, S.C. (2006). Rad52- mediated DNA annealing after Rad51-mediated DNA strand exchange promotes second ssDNA capture. EMBO J 25, 5539–5548.

59. Krasner, D.S., Daley, J.M., Sung, P., and Niu, H. (2015). Interplay between Ku and Replication Protein A in the Restriction of Exo1-mediated DNA Break End Resection. J. Biol. Chem. 290, 18806–18816.

60. Liu, L., Yan, Z., Osia, B.A., Twarowski, J., Sun, L., Kramara, J., Lee, R.S., Kumar, S., Elango, R., Li, H., et al. (2021). Tracking break-induced replication shows that it stalls at roadblocks. Nature 590, 655–659.

61. Theulot, B., Lacroix, L., Arbona, J.M., Millot, G.A., Jean, E., Cruaud, C., Pellet, J., Proux, F., Hennion, M., Engelen, S., et al. (2022). Genome-wide mapping of individual replication fork velocities using nanopore sequencing. Nat. Commun. 13, 3295.

62. Osman, F., Adriance, M., and McCready, S. (2000). The genetic control of spontaneous and UV-induced mitotic intrachromosomal recombination in the fission yeast *Schizosaccharomyces pombe*. Current genetics 38, 113–125.

63. Morrow, C.A., Nguyen, M.O., Fower, A., Wong, I.N., Osman, F., Bryer, C., and Whitby, M.C. (2017). Inter-Fork Strand Annealing causes genomic deletions during the termination of DNA replication. Elife 6. 10.7554/eLife.25490.

64. Goldstein, A.L., and McCusker, J.H. (1999). Three new dominant drug resistance cassettes for gene disruption in *Saccharomyces cerevisiae*. Yeast. 15, 1541–1553.

65. Ran, F.A., Hsu, P.D., Wright, J., Agarwala, V., Scott, D.A., and Zhang, F. (2013). Genome engineering using the CRISPR-Cas9 system. Nat. Protoc. 8, 2281–2308.

66. Basi, G., Schmid, E., and Maundrell, K. (1993). TATA box mutations in the Schizosaccharomyces pombe nmt1 promoter affect transcription efficiency but not the transcription start point or thiamine repressibility. Gene 123, 131–136.

67. Jacobs, J.Z., Ciccaglione, K.M., Tournier, V., and Zaratiegui, M. (2014). Implementation of the CRISPR-Cas9 system in fission yeast. Nat. Commun. 5, 5344.

68. Bahler, J., Wu, J.Q., Longtine, M.S., Shah, N.G., McKenzie, A., 3rd, Steever, A.B., Wach, A., Philippsen, P., and Pringle, J.R. (1998). Heterologous modules for efficient and versatile PCR-based gene targeting in Schizosaccharomyces pombe. Yeast 14, 943–951.

69. Maundrell, K. (1990). nmt1 of fission yeast. A highly transcribed gene completely repressed by thiamine. J. Biol. Chem. 265, 10857–10864.

70. Maundrell, K. (1993). Thiamine-repressible expression vectors pREP and pRIP for fission yeast. Gene. 123, 127–130.

71. Moreno, S., Klar, A., and Nurse, P. (1991). Molecular genetic analysis of fission yeast Schizosaccharomyces pombe. Methods Enzymol. 194, 795–823.

72. Osman, F., and Whitby, M.C. (2009). Monitoring homologous recombination following replication fork perturbation in the fission yeast *Schizosaccharomyces pombe*. Methods Mol. Biol. 521, 535–552.

