## Supplementary Material for "DNA nicks in both leading and lagging strand templates can trigger break-induced replication"

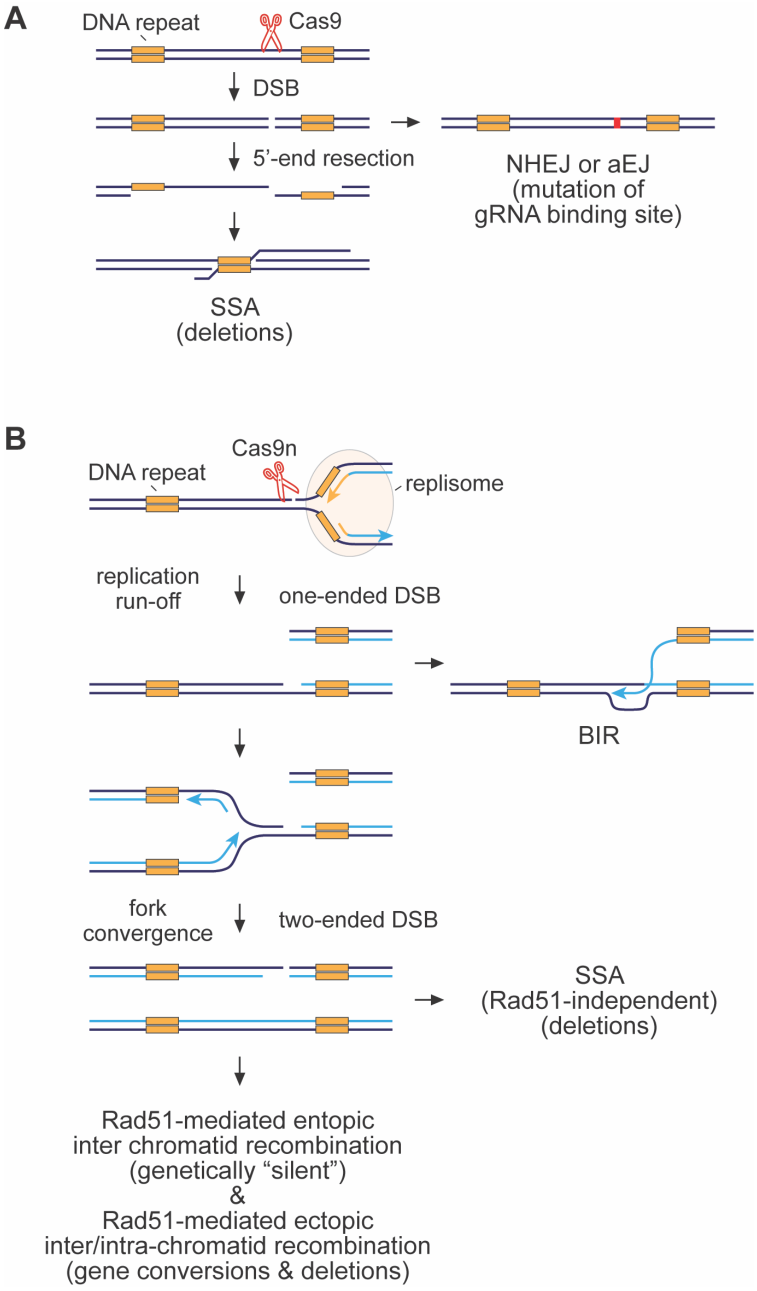

**Supplementary Figure 1. Models for the repair of DSBs induced by Cas9 and Cas9n**

**(A)** The repair of Cas9-induced DSBs. See main text for further details.

**(B)** The repair of DSBs caused by replication fork encounters with Cas9n-induced SSBs. See main text for further details.

### Supplementary Table 1: Recombination frequencies

<table>
<tr><th>Relevant Genotype and Strain no.</th><th>Cas9 variant or Flp-nick or GpII or RTS1</th><th>Position of reporter relative to SSB[a]</th><th>Number of colonies analysed (n)</th><th>Total Ade$^+$ recombinant frequency (x $10^{-4}$)<br>Median (95% CI)[b]</th><th>Ade$^+$ His$^+$ recombinant frequency (x $10^{-4}$)<br>gene conversions (GC)<br>Median (95% CI)[b]</th><th>Ade$^+$ His$^-$ recombinant frequency (x $10^{-4}$)<br>deletions (DEL)<br>Median (95% CI)[b]</th><th>P values[c]</th><th>Figure(s)</th></tr>
<tr><td>wild-type MCW9804<br>+ pLX3<br>(no gRNA)</td><td>no Cas9</td><td>flanking</td><td>11</td><td>2.800<br>(1.923 –6.232)</td><td>0.800<br>(0.3846 – 2.750)</td><td>2.308<br>(0.6061 – 4.638)</td><td></td><td>1D</td></tr>
<tr><td>wild-type MCW9804<br>+ pLX2<br>(+ gRNA)</td><td>no Cas9</td><td>flanking</td><td>12</td><td>1.950<br>(1.321 – 4.028)</td><td>0.4438<br>(0.3719 – 0.6250)</td><td>1.451<br>(0.9057 – 2.639)</td><td>total Ade+ vs MCW9804 + pLX3 total Ade+ = 0.0374<br>GC vs MCW9804 + pLX3 GC = 0.0335<br>DEL vs MCW9804 + pLX3 DEL = 0.3919</td><td>1D</td></tr>
<tr><td>wild-type MCW9806<br>+ pLX3<br>(no gRNA)</td><td>Cas9</td><td>flanking</td><td>35</td><td>2.667<br>(2.035 – 3.238)</td><td>0.6316<br>(0.4032 – 0.7463)</td><td>1.725<br>(1.482 – 2.000)</td><td></td><td>1D</td></tr>
<tr><td>wild-type MCW9806<br>+ pLX2<br>(+ gRNA)</td><td>Cas9</td><td>flanking</td><td>31</td><td>5482<br>(3535 – 6754)</td><td>64.10<br>(35.71 – 76.27)</td><td>5414<br>(3521 – 6717)</td><td>total Ade+ vs MCW9804 + pLX3 total Ade+ = <0.0001<br>GC vs MCW9806 + pLX3 GC = <0.0001<br>DEL vs MCW9806 + pLX3 DEL = <0.0001</td><td>1D</td></tr>
<tr><td>wild-type MCW10067<br>+ pLX3<br>(no gRNA)</td><td>Cas9d</td><td>flanking</td><td>26</td><td>2.006<br>(1.453 – 2.667)</td><td>0.4974<br>(0.3021 – 0.5935)</td><td>1.434<br>(1.086 – 1.901)</td><td></td><td>1D</td></tr>
<tr><td>wild-type MCW10067<br>+ pLX2<br>(+ gRNA)</td><td>Cas9d</td><td>flanking</td><td>26</td><td>5.190<br>(1.453 – 2.667)</td><td>2.273<br>(0.3021 – 0.5935)</td><td>2.784<br>(1.086 – 1.901)</td><td>total Ade+ vs MCW10067 + pLX3 total Ade+ = <0.0001<br>GC vs MCW10067 + pLX3 GC = <0.0001<br>DEL vs MCW10067 + pLX3 DEL = <0.0001</td><td>1D</td></tr>
<tr><td>wild-type MCW9831</td><td>Cas9n$^{H840A}$</td><td>flanking</td><td>26</td><td>3.153<br>(2.857 – 3.413)</td><td>0.6266<br>(0.5317 – 0.9148)</td><td>2.137<br>(1.595 – 2.778)</td><td></td><td>1D, 2C</td></tr>
</table>

| | | | | | | | | |
|---|---|---|---|---|---|---|---|---|
| + pLX3 (no gRNA) | | | | | | | | |
| wild-type MCW9831 + pLX2 (+ gRNA) | $Cas9n^{H840A}$ | flanking | 30 | 1066 (878.6 – 1174) | 85.78 (79.37 – 116.0) | 987.6 (783.6 – 1091) | GC vs MCW9806 + pLX2 GC = 0.0073<br>DEL vs MCW9806 + pLX2 DEL = <0.0001<br>total Ade+ vs MCW9831 + pLX3 total Ade+ = <0.0001<br>GC vs MCW9831 + pLX3 GC = <0.0001<br>DEL vs MCW9831 + pLX3 DEL = <0.0001 | 1D, 2C |
| wild-type MCW9834 + pLX3 (no gRNA) | $Cas9n^{D10A}$ | flanking | 27 | 2.681 (2.156 – 3.857) | 0.8803 (0.6579 – 1.262) | 1.462 (1.109 – 2.621) | | 1D, 2C |
| wild-type MCW9834 + pLX2 (+ gRNA) | $Cas9n^{D10A}$ | flanking | 27 | 717.0 (634.1 – 872.9) | 79.67 (56.34 – 115.2) | 624.2 (578.1 – 758.9) | GC vs MCW9806 + pLX2 GC = 0.1679<br>DEL vs MCW9806 +pLX2 DEL = <0.0001<br>total Ade+ vs MCW9834 + pLX3 total Ade+ = <0.0001<br>GC vs MCW9834 + pLX3 GC = <0.0001<br>DEL vs MCW9834 + pLX3 DEL = <0.0001 | 1D, 2C |
| *rad52*-R45A MCW10347 + pLX3 (no gRNA) | $Cas9n^{H840A}$ | flanking | 36 | 2.777 (2.308 – 3.660) | 0.3856 (0.3344 – 0.4720) | 1.989 (1.783 – 3.500) | | 2C |
| *rad52*-R45A MCW10347 + pLX2 (+ gRNA) | $Cas9n^{H840A}$ | flanking | 36 | 565.9 (517.2 – 634.6) | 77.68 (69.51 – 81.34) | 487.1 (420.1 – 557.9) | GC vs MCW9831 + pLX2 GC = 0.4272<br>DEL vs MCW9831 +pLX2 DEL = <0.0001 | 2C |

| | | | | | | | | |
|---|---|---|---|---|---|---|---|---|
| *rad52*-R45A MCW10343<br>+ pLX3<br>(no gRNA) | Cas9n$^{D10A}$ | flanking | 34 | 3.788 (2.215 – 5.703) | 0.6108 (0.5024 – 0.7277) | 2.441 (1.489 – 4.836) | | 2C |
| *rad52*-R45A MCW10343<br>+ pLX2<br>(+ gRNA) | Cas9n$^{D10A}$ | flanking | 35 | 496.1 (444.6 – 585.5) | 85.24 (73.58 – 93.41) | 415.5 (353.8 – 484.2) | GC vs MCW9834 + pLX2 GC = >0.9999<br>DEL vs MCW9834 +pLX2 DEL = 0.0029 | 2C |
| *rad51*Δ MCW10333<br>+ pLX3<br>(no gRNA) | Cas9n$^{H840A}$ | flanking | 24 | 22.10 (17.79 – 31.61) | 0.1241 (0.1000 – 0.1748) | 21.85 (17.71 – 31.50) | | 2C |
| *rad51*Δ MCW10333<br>+ pLX2<br>(+ gRNA) | Cas9n$^{H840A}$ | flanking | 23 | 3538 (2952 – 4237) | 9.302 (6.438 – 11.74) | 3531 (2942 – 4228) | GC vs MCW9831 + pLX2 GC = <0.0001<br>DEL vs MCW9831 +pLX2 DEL = 0.0002 | 2C |
| *rad51*Δ MCW10330<br>+ pLX3<br>(no gRNA) | Cas9n$^{D10A}$ | flanking | 24 | 9.675 (5.419 – 16.40) | 0.1033 (0.0713 – 0.1493) | 9.487 (5.302 – 16.35) | | 2C |
| *rad51*Δ MCW10330<br>+ pLX2<br>(+ gRNA) | Cas9n$^{D10A}$ | flanking | 24 | 3602 (3412 – 4250) | 11.61 (8.772 – 15.00) | 3587 (3404 – 4238) | GC vs MCW9834 + pLX2 GC = <0.0001<br>DEL vs MCW9834 +pLX2 DEL = <0.0001 | 2C |
| *rad51*Δ *rad52*Δ MCW10342<br>+ pLX3<br>(no gRNA) | Cas9n$^{H840A}$ | flanking | 15 | 0.8578 (0.6068 – 1.121) | 0.0196 (0.0000 – 0.02439) | 0.8578 (0.5825 – 1.121) | | 2C |
| *rad51*Δ *rad52*Δ MCW10342<br>+ pLX2<br>(+ gRNA) | Cas9n$^{H840A}$ | flanking | 15 | 769.3 (602.5 – 880.7) | 1.731 (0.9346 – 2.113) | 768.1 (601.8 – 878.4) | GC vs MCW9831 + pLX2 GC = <0.0001<br>DEL vs MCW9831 +pLX2 DEL = 0.6405<br>GC vs MCW10333 + pLX2 GC = <0.0001<br>DEL vs MCW10333 +pLX2 DEL = <0.0001 | 2C |

| | | | | | | | | |
|---|---|---|---|---|---|---|---|---|
| *rad51*Δ *rad52*Δ MCW10335 + pLX3 (no gRNA) | Cas9n$^{D10A}$ | flanking | 13 | 1.5 (0.9231 – 4.492) | 0.1033 (0.000 – 0.06557) | 9.487 (0.9205 – 4.426) | | 2C |
| *rad51*Δ *rad52*Δ MCW10335 + pLX2 (+ gRNA) | Cas9n$^{D10A}$ | flanking | 24 | 1167 (667.1 – 1476) | 1.208 (1.053 – 1.389) | 1166 (666.3 – 1474) | GC vs MCW9834 + pLX2 GC = <0.0001 DEL vs MCW9834 +pLX2 DEL = 0.1479 GC vs MCW10330 + pLX2 GC = <0.0001 DEL vs MCW10330 +pLX2 DEL = <0.0001 | 2C |
| wild-type MCW7229 + pLX3 (no gRNA) | no Cas9 | 0.7 kb down-stream | 30 | 2.437 (2.186 – 3.715) | 0.9568 (0.7979 – 1.246) | 1.829 (1.285 – 2.409) | | 3B |
| wild-type MCW7229 + pLX32 (+ gRNA) | no Cas9 | 0.7 kb down-stream | 29 | 2.459 (1.977 – 3.705) | 0.8466 (0.6119 – 1.011) | 1.535 (1.127 – 2.784) | total Ade+ vs MCW7229 + pLX3 total Ade+ = 0.69 | 3B |
| wild-type MCW10229 + pLX3 (no gRNA) | Cas9 | 0.7 kb down-stream | 14 | 1.960 (1.601 – 3.185) | 0.6598 (0.4825 – 0.8635) | 1.435 (1.111 – 2.332) | | 3B |
| wild-type MCW10229 + pLX32 (+ gRNA) | Cas9 | 0.7 kb down-stream | 14 | 2.543 (1.406 – 4.167) | 0.6737 (0.4219 – 0.9820) | 2.196 (0.7884 – 3.086) | total Ade+ vs MCW10229 + pLX3 total Ade+ = 0.7345 | 3B |
| wild-type MCW10072 + pLX3 (no gRNA) | Cas9d | 0.7 kb down-stream | 26 | 2.207 (1.775 – 3.452) | 0.7349 (0.5230 – 1.031) | 1.340 (1.034 – 1.976) | | 3B |
| wild-type MCW10072 + pLX32 (+ gRNA) | Cas9d | 0.7 kb down-stream | 21 | 2.227 (1.873 – 3.322) | 0.65 (0.5272 – 0.9140) | 1.445 (1.237 – 2.066) | total Ade+ vs MCW10072 + pLX3 total Ade+ = 0.5597 | 3B |
| wild-type MCW10236 + pLX3 (no gRNA) | Cas9n$^{H840A}$ | 0.7 kb down-stream | 27 | 2.507 (1.560 – 3.384) | 0.6884 (0.4926 – 1.262) | 1.186 (0.9697 – 2.150) | | 3B |
| wild-type MCW10236 + pLX32 (+ gRNA) | Cas9n$^{H840A}$ | 0.7 kb down-stream | 26 | 34.06 (30.25 – 47.30) | 3.420 (2.182 – 4.198) | 30.43 (27.05 – 42.29) | total Ade+ vs MCW10236 + pLX3 total Ade+ = <0.0001 | 3B |

| | | | | | | | | |
|---|---|---|---|---|---|---|---|---|
| wild-type MCW10232 + pLX3 (no gRNA) | Cas9n$^{D10A}$ | 0.7 kb down-stream | 26 | 2.164 (1.685 – 2.654) | 0.6695 (0.4134 – 0.9804) | 1.370 (1.077 – 1.857) | | 3B |
| wild-type MCW10232 + pLX32 (+ gRNA) | Cas9n$^{D10A}$ | 0.7 kb down-stream | 27 | 122.3 (47.71 – 157.6) | 12.01 (6.838 – 17.77) | 105.6 (29.52 – 135.1) | total Ade+ vs MCW10232 + pLX3 total Ade+ = <0.0001 | 3B |
| wild-type MCW10178 + pLX47 (no gRNA) | Cas9n$^{H840A}$ | 13.8 kb down-stream | 48 | 3.264 (2.592 – 3.855) | 0.6974 (0.5310 – 1.093) | 1.950 (1.629 – 2.710) | | 3C |
| wild-type MCW10178 + pLX46 (+ gRNA) | Cas9n$^{H840A}$ | 13.8 kb down-stream | 48 | 3.881 (3.409 – 4.348) | 0.8578 (0.7143 – 1.042) | 2.788 (2.308 – 3.188) | total Ade+ vs MCW10178 + pLX47 total Ade+ = 0.009 | 3C |
| wild-type MCW10174 + pLX47 (no gRNA) | Cas9n$^{D10A}$ | 13.8 kb down-stream | 35 | 2.484 (2.222 – 2.733) | 0.6897 (0.5580 – 0.7813) | 1.852 (1.518 – 2.010) | | 3C |
| wild-type MCW10174 + pLX46 (+ gRNA) | Cas9n$^{D10A}$ | 13.8 kb down-stream | 34 | 5.025 (4.633 – 6.335) | 1.242 (0.9828 – 1.512) | 3.786 (3.235 – 4.483) | total Ade+ vs MCW10174 + pLX47 total Ade+ = <0.0001 | 3C |
| *ori-1253*Δ MCW10481 + pLX47 (no gRNA) | Cas9n$^{H840A}$ | 13.8 kb down-stream | 43 | 5.514 (4.980 – 6.354) | 1.215 (0.9804 – 1.508) | 4.181 (3.398 – 4.548) | | 3C, 4C |
| *ori-1253*Δ MCW10481 + pLX46 (+ gRNA) | Cas9n$^{H840A}$ | 13.8 kb down-stream | 47 | 18.38 (15.43 – 21.17) | 1.848 (1.310 – 2.389) | 15.20 (12.74 – 17.94) | total Ade+ vs MCW10481 + pLX47 total Ade+ = <0.0001 | 3C, 4C |
| *ori-1253*Δ MCW10473 + pLX47 (no gRNA) | Cas9n$^{D10A}$ | 13.8 kb down-stream | 33 | 5.288 (5.030 – 5.761) | 0.9124 (0.7991 – 1.154) | 4.240 (3.700 – 4.767) | | 3C, 4C |
| *ori-1253*Δ MCW10473 + pLX46 (+ gRNA) | Cas9n$^{D10A}$ | 13.8 kb down-stream | 36 | 18.96 (15.52 – 20.93) | 1.641 (1.310 – 2.581) | 15.86 (13.90 – 19.09) | total Ade+ vs MCW10473 + pLX47 total Ade+ = <0.0001 | 3C, 4C |
| *ori-1253*Δ *ku70*Δ MCW10708 + pLX47 (no gRNA) | Cas9n$^{H840A}$ | 13.8 kb down-stream | 45 | 7.538 (6.861 – 8.909) | 1.269 (0.9804 – 1.581) | 6.082 (5.729 – 7.333) | | 3C |
| *ori-1253*Δ *ku70*Δ MCW10708 + pLX46 | Cas9n$^{H840A}$ | 13.8 kb down-stream | 48 | 43.61 (37.42 – 47.27) | 3.745 (3.371 – 4.785) | 36.81 (31.55 – 40.81) | total Ade+ vs MCW10708 + pLX47 total Ade+ = <0.0001 | 3C |

| | | | | | | | | |
|---|---|---|---|---|---|---|---|---|
| (+ gRNA) | | | | | | | total Ade+ vs MCW10481 + pLX46 total Ade+ = <0.0001 | |
| *ori-1253Δ ku70Δ* MCW10706 + pLX47 (no gRNA) | Cas9n$^{D10A}$ | 13.8 kb down-stream | 24 | 7.068 (6.317 – 8.715) | 1.332 (0.9600 – 1.710) | 5.953 (5.060 – 7.346) | | 3C |
| *ori-1253Δ ku70Δ* MCW10706 + pLX46 (+ gRNA) | Cas9n$^{D10A}$ | 13.8 kb down-stream | 23 | 46.22 (42.45 – 50.24) | 3.198 (2.620 – 5.333) | 41.11 (36.16 – 44.27) | total Ade+ vs MCW10706 + pLX47 total Ade+ = <0.0001 total Ade+ vs MCW10473 + pLX46 total Ade+ = <0.0001 | 3C |
| wild-type MCW9835 + pLX3 (no gRNA) | Cas9n$^{H840A}$ | flanking* | 17 | 2.653 (2.191 – 3.353) | 0.6596 (0.5666 – 1.032) | 1.519 (1.364 – 2.099) | | 4B |
| wild-type MCW9835 + pLX2 (+ gRNA) | Cas9n$^{H840A}$ | flanking* | 18 | 1524 (1418 – 1856) | 21.42 (20.20 – 23.32) | 1501 (1398 – 1835) | total Ade+ vs MCW9835 + pLX3 total Ade+ = <0.0001 | 4B |
| wild-type MCW9525 + pLX3 (no gRNA) | Cas9n$^{D10A}$ | flanking* | 14 | 2.451 (1.854 – 3.351) | 0.5670 (0.4839 – 0.8537) | 1.682 (1.340 – 2.022) | | 4B |
| wild-type MCW9525 + pLX2 (+ gRNA) | Cas9n$^{D10A}$ | flanking* | 15 | 1069 (954.2 – 1285) | 426.5 (391.7 – 449.7) | 679.5 (557.9 – 857.9) | total Ade+ vs MCW9525 + pLX3 total Ade+ = <0.0001 | 4B |
| wild-type MCW10107 + pREP1 | no Flp$^{H305L}$ TS *FRT* | flanking* | 14 | 3.065 (1.833 – 5.133) | 0.4750 (0.2597 – 0.7725) | 2.574 (1.515 – 3.991) | | 4B |
| wild-type MCW10107 + pREP1-Flp$^{H305L}$ | Flp$^{H305L}$ TS *FRT* | flanking* | 15 | 817.4 (744.6 – 890.1) | 87.21 (76.64 – 115.9) | 704.4 (640.7 – 843.4) | total Ade+ vs MCW10107 + pREP1 total Ade+ = <0.0001 | 4B |
| wild-type MCW10108 + pREP1 | no Flp$^{H305L}$ BS *FRT* | flanking* | 15 | 3.032 (1.939 – 3.352) | 0.5051 (0.3226 – 1.194) | 1.908 (1.576 – 2.772) | | 4B |
| wild-type MCW10108 + pREP1-Flp$^{H305L}$ | Flp$^{H305L}$ BS *FRT* | flanking* | 15 | 1112 (811.6 – 1311) | 73.39 (57.97 – 80.75) | 1038 (757.0 – 1211) | total Ade+ vs MCW10108 + pREP1 total Ade+ = <0.0001 | 4B |
| wild-type JSA267 + pREP1 | no GpII | flanking* | 21 | 2.727 (1.650 – 3.594) | 0.3646 (0.2922 – 0.7009) | 2.037 (1.359 – 2.673) | | 4B |

| | | | | | | | | |
|---|---|---|---|---|---|---|---|---|
| | TS GpII cleavage site | | | | | | | |
| wild-type JSA267<br>+ pREP1-GpII | GpII<br>TS GpII cleavage site | flanking* | 23 | 984.0<br>(735.8 – 1118) | 40.91<br>(37.74 – 64.52) | 914.9<br>(698.1 – 1077) | total Ade+ vs JSA267 + pREP1 total Ade+ = <0.0001 | 4B |
| wild-type MCW1159<br>+ pREP1 | no GpII<br>BS GpII cleavage site | flanking* | 23 | 2.103<br>(1.712 – 3.111) | 0.4455<br>(0.3571 – 0.5828) | 1.714<br>(1.332 – 2.398) | | 4B |
| wild-type MCW1159<br>+ pREP1-GpII | GpII<br>BS GpII cleavage site | flanking* | 23 | 1132<br>(1005 – 1230) | 40.91<br>(27.56 – 63.03) | 1077<br>(958.7 – 1165) | total Ade+ vs MCW1159 + pREP1 total Ade+ = <0.0001 | 4B |
| *ori-1253Δ* MCW10798<br>+ pREP1 | no $Flp^{H305L}$<br>TS *FRT* | 13.4 kb down-stream | 43 | 3.901<br>(3.421 – 4.923) | 0.8198<br>(0.5839 – 0.9419) | 2.805<br>(2.121 – 3.389) | | 4C |
| *ori-1253Δ* MCW10798<br>+ pREP1-$Flp^{H305L}$ | $Flp^{H305L}$<br>TS *FRT* | 13.4 kb down-stream | 45 | 61.34<br>(53.73 – 72.90) | 3.874<br>(3.590 – 4.202) | 57.39<br>(49.40 – 69.27) | total Ade+ vs MCW10798 + pREP1 total Ade+ = <0.0001<br>total Ade+ vs MCW10481 + pLX46 total Ade+ = <0.0001<br>total Ade+ vs MCW10473 + pLX46 total Ade+ = <0.0001 | 4C |
| *ori-1253Δ* MCW10853<br>+ pREP1 | no $Flp^{H305L}$<br>BS *FRT* | 13.4 kb down-stream | 16 | 6.518<br>(4.844 – 7.964) | 1.162<br>(0.8203 – 1.696) | 5.231<br>(3.691 – 6.674) | | 4C |
| *ori-1253Δ* MCW10853<br>+ pREP1-$Flp^{H305L}$ | $Flp^{H305L}$<br>BS *FRT* | 13.4 kb down-stream | 23 | 61.29<br>(47.09 – 98.95) | 4.977<br>(2.913 – 7.512) | 50.86<br>(41.26 – 95.79) | total Ade+ vs MCW10853 + pREP1 total Ade+ = <0.0001<br>total Ade+ vs MCW10481 + pLX46 total Ade+ = <0.0001<br>total Ade+ vs MCW10473 + pLX46 total Ade+ = <0.0001 | 4C |
| *ori-1253Δ* MCW10190<br>+ pREP1 | no GpII<br>TS GpII cleavage site | 13.4 kb down-stream | 43 | 5.956<br>(4.967 – 6.709) | 1.783<br>(1.330 – 2.077) | 3.792<br>(2.959 – 4.910) | | 4C |

| | | | | | | | | |
|---|---|---|---|---|---|---|---|---|
| *ori-1253*Δ MCW10190<br><br>+ pREP1-GpII | GpII<br><br>TS GpII cleavage site | 13.4 kb down-stream | 44 | 42.09 (36.50 – 55.05) | 4.351 (3.375 – 5.000) | 35.09 (32.39 – 47.22) | total Ade+ vs MCW10190 + pREP1 total Ade+ = <0.0001<br><br>total Ade+ vs MCW10481 + pLX46 total Ade+ = <0.0001<br><br>total Ade+ vs MCW10473 + pLX46 total Ade+ = <0.0001 | 4C |
| *ori-1253*Δ MCW10194<br><br>+ pREP1 | no GpII<br><br>BS GpII cleavage site | 13.4 kb down-stream | 43 | 5.313 (4.420 – 6.196) | 1.953 (1.635 – 2.110) | 3.034 (2.431 – 3.840) | | 4C |
| *ori-1253*Δ MCW10194<br><br>+ pREP1-GpII | GpII<br><br>BS GpII cleavage site | 13.4 kb down-stream | 45 | 72 (62.20 – 93.04) | 12.53 (10.26 – 14.53) | 53.5 (47.90 – 63.49) | total Ade+ vs MCW10194 + pREP1 total Ade+ = <0.0001<br><br>total Ade+ vs MCW10481 + pLX46 total Ade+ = <0.0001<br><br>total Ade+ vs MCW10473 + pLX46 total Ade+ = <0.0001 | 4C |

[a] The different recombination reporters are shown in Figure 1A and 3A.
[b] The values in parentheses are the 95% confidence interval.
[c] Details of the statistical analysis are in Supplementary Data 1

**Supplementary Table 2:** ***Schizosaccharomyces pombe*** **strains** (in order of appearance)

| **Strain No.** | **Relevant genotype** | **Source** |
|---|---|---|
| MCW9804 | $h^+$ *ade6-M375* int::*pUC8/his3+/2kb spacer/ade6-L469 ura4-D8 leu1-32 his3-D1 arg3-D4* | This study |
| MCW9806 | $h^+$ *lys1*::P*nmt81-cas9-hphMX4 ade6-M375* int::*pUC8/his3+/2kb spacer/ade6-L469 ura4-D8 leu1-32 his3-D1 arg3-D4* | This study |
| MCW10067 | $h^-$ *lys1*::P*nmt81-cas9d-hphMX4 ade6-M375* int::*pUC8/his3+/2kb spacer/ade6-L469 ura4-D8 leu1-32 his3-D1 arg3-D4* | This study |
| MCW9831 | $h^+$ *lys1*::P*nmt81-cas9nH840A-hphMX4 ade6-M375* int::*pUC8/his3+/2kb spacer/ade6-L469 ura4-D8 leu1-32 his3-D1 arg3-D4* | This study |
| MCW9834 | $h^+$ *lys1*::P*nmt81-cas9nD10A-hphMX4 ade6-M375* int::*pUC8/his3+/2kb spacer/ade6-L469 ura4-D8 leu1-32 his3-D1 arg3-D4* | This study |
| MCW10347 | $h^+$ *rad52Δ::rad52+-kanMX6 lys1*::P*nmt81-cas9nH840A-hphMX4 ade6-M375* int::*pUC8/his3+/2kb spacer/ade6-L469 ura4-D8 leu1-32 his3-D1 arg3-D4* | This study |
| MCW10343 | $h^+$ *rad52Δ::rad52+-kanMX6 lys1*::P*nmt81-cas9nD10A-hphMX4 ade6-M375* int::*pUC8/his3+/2kb spacer/ade6-L469 ura4-D8 leu1-32 his3-D1 arg3-D4* | This study |
| MCW10333 | $h^+$ *rad51Δ::arg3+ lys1*::P*nmt81-cas9nH840A-hphMX4 ade6-M375* int::*pUC8/his3+/2kb spacer/ade6-L469 ura4-D8 leu1-32 his3-D1 arg3-D4* | This study |
| MCW10330 | $h^-$ *rad51Δ::arg3+ lys1*::P*nmt81-cas9nD10A-hphMX4 ade6-M375* int::*pUC8/his3+/2kb spacer/ade6-L469 ura4-D8 leu1-32 his3-D1 arg3-D4* | This study |
| MCW10342 | $h^+$ *rad51Δ::arg3+ rad52Δ::kanMX6 lys1*::P*nmt81-cas9nH840A-hphMX4 ade6-M375* int::*pUC8/his3+/2kb spacer/ade6-L469 ura4-D8 leu1-32 his3-D1 arg3-D4* | This study |
| MCW10335 | $h^-$ *rad51Δ::arg3+ rad52Δ::kanMX6 lys1*::P*nmt81-cas9nD10A-hphMX4 ade6-M375* int::*pUC8/his3+/2kb spacer/ade6-L469 ura4-D8 leu1-32 his3-D1 arg3-D4* | This study |
| MCW7229 | $h^+$ (12.4 kb from *ade6*)int::*ade6-M375 int::pUC8/*$his3^+$*/ade6-L469/kanMX6 ade6-D1 ura4-D18 his3-D1 leu1-32 arg3-D4* | Jalan et al 2019 |
| MCW10229 | $h^+$ *lys1*::P*nmt81-cas9-hphMX4* (12.4 kb from *ade6*)int::*ade6-M375 int::pUC8/*$his3^+$*/ade6-L469/kanMX6 ade6-D1 ura4-D18 his3-D1 leu1-32 arg3-D4* | This study |
| MCW10072 | $h^+$ *lys1*::P*nmt81-cas9d-hphMX4* (12.4 kb from *ade6*)int::*ade6-M375 int::pUC8/*$his3^+$*/ade6-L469/kanMX6 ade6-D1 ura4-D18 his3-D1 leu1-32 arg3-D4* | This study |
| MCW10236 | $h^-$ *lys1*::P*nmt81-cas9nH840A-hphMX4* (12.4 kb from *ade6*)int::*ade6-M375 int::pUC8/*$his3^+$*/ade6-L469/kanMX6 ade6-D1 ura4-D18 his3-D1 leu1-32 arg3-D4* | This study |
| MCW10232 | $h^-$ *lys1*::P*nmt81-cas9nD10A-hphMX4* (12.4 kb from *ade6*)int::*ade6-M375 int::pUC8/*$his3^+$*/ade6-L469/kanMX6 ade6-D1 ura4-D18 his3-D1 leu1-32 arg3-D4* | This study |
| MCW10178 | $h^+$ *lys1*::P*nmt81-cas9nH840A-hphMX4* (12.4 kb from *ade6*)int::*ade6-M375 int::pUC8/*$his3^+$*/ade6-L469/ura4MX4 ade6Δ::kanMX6 ura4-D18 his3-D1 leu1-32 arg3-D4* | This study |
| MCW10174 | $h^+$ *lys1*::P*nmt81-cas9nD10A-hphMX4* (12.4 kb from *ade6*)int::*ade6-M375 int::pUC8/*$his3^+$*/ade6-L469/ura4MX4 ade6Δ::kanMX6 ura4-D18 his3-D1 leu1-32 arg3-D4* | This study |
| MCW10481 | $h^-$ *oriIII-1253Δ::natMX4 lys1*::P*nmt81-cas9nH840A-hphMX4* (12.4 kb from *ade6*)int::*ade6-M375 int::pUC8/*$his3^+$*/ade6-L469/ura4MX4 ade6Δ::kanMX6 ura4-D18 his3-D1 leu1-32 arg3-D4* | This study |
| MCW10473 | $h^+$ *oriIII-1253Δ::natMX4 lys1*::P*nmt81-cas9nD10A-hphMX4* (12.4 kb from *ade6*)int::*ade6-M375 int::pUC8/*$his3^+$*/ade6-L469/ura4MX4 ade6Δ::kanMX6 ura4-D18 his3-D1 leu1-32 arg3-D4* | This study |
| MCW10708 | $h^+$ *ku70Δ::arg3MX4 oriIII-1253Δ::natMX4 lys1*::P*nmt81-cas9nH840A-hphMX4* (12.4 kb from *ade6*)int::*ade6-M375 int::pUC8/*$his3^+$*/ade6-L469/ura4MX4 ade6Δ::kanMX6 ura4-D18 his3-D1 leu1-32 arg3-D4* | This study |
| MCW10706 | $h^-$ *ku70Δ::arg3MX4 oriIII-1253Δ::natMX4 lys1*::P*nmt81-cas9nD10A-hphMX4* (12.4 kb from *ade6*)int::*ade6-M375 int::pUC8/*$his3^+$*/ade6-L469/ura4MX4 ade6Δ::kanMX6 ura4-D18 his3-D1 leu1-32 arg3-D4* | This study |
| MCW9835 | $h^+$ *lys1*::P*nmt81-cas9nH840A-hphMX4 ade6-M375* int::*pUC8/his3+/ade6-L469 ura4-D8 leu1-32 his3-D1 arg3-D4* | This study |
| MCW9525 | $h^+$ *lys1*::P*nmt81-cas9nD10A-hphMX4 ade6-M375* int::*pUC8/his3+/ade6-L469 ura4-D8 leu1-32 his3-D1 arg3-D4* | This study |

| | | |
|---|---|---|
| MCW10107 | *h⁻ ade6-M375* int::*pUC8*/*his3+*/*FRT* TS/*ade6-L469 ura4-D8 leu1-32 his3-D1 arg3-D4* | This study |
| MCW10108 | *h⁻ ade6-M375* int::*pUC8*/*his3+*/*FRT* BS/*ade6-L469 ura4-D8 leu1-32 his3-D1 arg3-D4* | This study |
| JSA267 | *h⁻ ade6-M375* int::*pUC8*/*his3+*/gpII cleavage site TS/*ade6-L469 ura4-D8 leu1-32 his3-D1* | Osman et al 2016 |
| MCW1159 | *h⁻ ade6-M375* int::*pUC8*/*his3+*/gpII cleavage site BS/*ade6-L469 ura4-D8 leu1-32 his3-D1* | Osman et al 2016 |
| MCW10798 | *h⁻ oriIII-1253*Δ::*natMX4* (12.4 kb from *ade6*)int::*ade6-M375 int*::*pUC8/his3⁺/ade6-L469/ura4MX4 ade6*Δ::*FRT* TS-*hphMX4 ura4-D18 his3-D1 leu1-32 arg3-D4* | This study |
| MCW10853 | *h⁻ oriIII-1253*Δ::*natMX4* (12.4 kb from *ade6*)int::*ade6-M375 int*::*pUC8/his3⁺/ade6-L469/ura4MX4 ade6*Δ::*FRT* BS-*hphMX4 ura4-D18 his3-D1 leu1-32 arg3-D4* | This study |
| MCW10190 | *h⁺ oriIII-1253*Δ::*natMX4* (12.4 kb from *ade6*)int::*ade6-M375 int*::*pUC8/his3⁺/ade6-L469/ura4MX4 ade6*Δ::gpII cleavage site TS-*hphMX4 ura4-D18 his3-D1 leu1-32 arg3-D4* | This study |
| MCW10194 | *h⁺ oriIII-1253*Δ::*natMX4* (12.4 kb from *ade6*)int::*ade6-M375 int*::*pUC8/his3⁺/ade6-L469/ura4MX4 ade6*Δ::gpII cleavage site BS-*hphMX4 ura4-D18 his3-D1 leu1-32 arg3-D4* | This study |
